# Decoding the architecture of the varicella-zoster virus transcriptome

**DOI:** 10.1101/2020.05.25.110965

**Authors:** Shirley E. Braspenning, Tomohiko Sadaoka, Judith Breuer, Georges M.G.M Verjans, Werner J.D. Ouwendijk, Daniel P. Depledge

**Author notes:** co-senior authors (Werner J.D. Ouwendijk, Daniel P. Depledge).

## Abstract

Varicella-zoster virus (VZV), a double-stranded DNA virus, causes varicella, establishes lifelong latency in ganglionic neurons, and reactivates later in life to cause herpes zoster, commonly associated with chronic pain. The VZV genome is densely packed and produces multitudes of overlapping transcripts deriving from both strands. While 71 distinct open reading frames (ORFs) have thus far been experimentally defined, the full coding potential of VZV remains unknown. Here, we integrated multiple short-read RNA sequencing approaches with long-read direct RNA sequencing on RNA isolated from VZV-infected cells to provide a comprehensive reannotation of the lytic VZV transcriptome architecture. Through precise mapping of transcription start sites, splice junctions, and polyadenylation sites, we identified 136 distinct polyadenylated VZV RNAs that encode canonical ORFs, non-canonical ORFs, and ORF fusions, as well as putative non-coding RNAs (ncRNAs). Furthermore, we determined the kinetic class of all VZV transcripts and observed, unexpectedly, that transcripts encoding the ORF62 protein, previously designated as *immediate-early,* were expressed with *late* kinetics. Our work showcases the complexity of the VZV transcriptome and provides a comprehensive resource that will facilitate future functional studies of coding RNAs, ncRNAs, and the biological mechanisms underlying the regulation of viral transcription and translation during lytic VZV infection.

## Introduction

Varicella-zoster virus (VZV) is a ubiquitous human alphaherpesvirus and causative agent of both varicella (chickenpox) and herpes zoster (HZ or shingles) (Gershon et al., 2015). Varicella results from primary VZV infection and leads to the establishment of a lifelong latent infection in sensory neurons of the trigeminal and dorsal root ganglia (Depledge et al., 2018a; Gilden et al., 1983). In one-third of infected individuals, VZV reactivates from latency later in life to cause HZ (Gershon et al., 2015). Whereas varicella is generally experienced as a benign childhood disease, HZ is frequently associated with difficult-to-treat chronic pain (post-herpetic neuralgia) (Gilden et al., 2000; Johnson and Rice, 2014). Despite the recent availability of the highly effective HZ subunit vaccine (Shingrix), the health and societal burden of HZ and its complications remains high due to adverse side effects, the high cost of the vaccines, and changing demographics (Gater et al., 2015).

The 125 kb double-stranded DNA (dsDNA) genome of VZV, first sequenced in 1986, encodes at least 71 unique open-reading frames (ORFs) that are expressed during lytic infection (Cohen, 2010; Davison and Scott, 1986). The current annotation of the VZV genome largely relies on both *in silico* ORF predictions and homologous ORFs in the closely related human alphaherpesvirus herpes simplex virus type 1 (HSV-1) (Cohen, 2010). For most VZV ORFs, the boundaries of transcription are not accurately determined meaning that transcription start sites and polyadenylation sites are poorly resolved. Moreover, *in silico* ORF prediction does not account for the possibility of spliced transcripts. Indeed, we recently applied ultra-deep short-read RNA-sequencing to define the latent viral transcriptome and discovered the spliced VZV latency-associated transcript (VLT) (Depledge et al., 2018b).

The compact nature of viral genomes, combined with their ability to encode overlapping RNAs, presents a significant challenge to studies that rely on interrogating genome sequences or that use viral mutants to probe the function(s) of viral proteins. Here, a single point mutation or frameshift may impact multiple distinct RNAs at the same time, a factor that may be further confounded by inaccurate or missing transcript annotations. The re-annotation of the human cytomegalovirus (HCMV) transcriptional and translational landscape and subsequent refinements of HSV-1, Kaposi’s sarcoma-associated herpesvirus (KSHV), and human herpesvirus 6 (HHV6) transcriptome architectures have all demonstrated that herpesviruses exhibit a complex transcriptional pattern of alternative splicing, opposing transcription, read-through transcription, fusion transcripts, 5’ untranslated region (5’ UTR) and 3’ UTR variations and previously unidentified non-coding RNAs (Arias et al., 2014; Bencun et al., 2018; Depledge et al., 2019; Finkel et al., 2020; O’Grady et al., 2016, 2019; Stern-Ginossar et al., 2012; Whisnant et al., 2019). Indeed, recent cDNA-based long-read sequencing also indicated that the lytic VZV transcriptome is substantially more complex than previously recognized (Prazsák et al., 2018).

By analogy to other herpesviruses and limited experimental data (Lenac Rovis et al., 2013; Reichelt et al., 2009), VZV transcripts and their encoded proteins have been divided into three kinetic classes: immediate-early (*IE*), early (*E*) and late (*L*). Expression of *E* and *L* transcripts is considered dependent on viral proteins of the preceding kinetic classes, while expression of *IE* transcripts occurs in the absence of viral protein synthesis (Honess and Roizman, 1974). Prior studies have defined four VZV proteins encoded by ORF4, ORF61, ORF62, and ORF63 as being transcriptional regulators that initiate lytic transcript expression (Defechereux et al., 1993; Kost et al., 1995; Moriuchi et al., 1993, 1994; Perera et al., 1993), whose corresponding transcripts have been classified as *IE* by analogy to their HSV-1 orthologues. *L* transcripts, such as VLTly, the lytic isoform of VLT, are either expressed at very low levels prior to or exclusively after viral DNA replication has commenced (Depledge et al., 2018b). However, the species specificity and highly cell-associated nature of VZV *in vitro* have hampered detailed analysis of VZV transcription. Improved protocols to obtain cell-free VZV and mass-spectrometry have provided some insight into the temporal pattern of viral protein expression (Ouwendijk et al., 2020), but lack sensitivity compared to RNA-sequencing and do not provide information on viral transcription.

To address this, we have decoded the architecture of the lytic VZV transcriptome in both human epithelial cells and neurons while contrasting discrete VZV strains. We subsequently established the kinetic class of all lytic viral transcripts and integrated these results to provide a comprehensive overview of the complexity and structure of the lytic VZV transcriptome as a rich resource that will enhance future functional studies of VZV biology.

## Results

### Decoding the complexity of lytic VZV gene expression

Standard methods for annotating viral transcriptomes require the integration of multiple types of Illumina RNA sequencing (RNA-Seq) data to identify transcription start sites (TSS), cleavage and polyadenylation sites (CPAS), splice sites, and transcript structures. The latter is particularly challenging to infer using conventional short-read sequencing approaches (Depledge et al., 2018c). By contrast, direct RNA sequencing (dRNA-Seq) using nanopore arrays offers the potential to capture all these distinct data points in a single sequencing run (Depledge et al., 2019; Garalde et al., 2018). To examine the structure of the lytic VZV transcriptome, ARPE-19 cells were infected with the VZV pOka wild-type strain and total RNA was extracted at 96 hours post infection (hpi, Figure 1A). The 96 hpi time-point was chosen to maximise the diversity of VZV transcripts likely to be present. Sequencing of the polyadenylated RNA fraction was performed using both RNA-Seq and dRNA-Seq (Figure 1A). Whereas short reads generated by standard Illumina RNA-Seq are not amenable for accurate isoform reconstruction in complex reads, they provide higher sequencing depth, detection of CPAS, and, crucially, enables splice-site correction of dRNA-Seq reads via the junction-polishing package in the software FLAIR (Tang et al., 2020). By contrast, dRNA-Seq can sequence full-length RNAs and provides critical information on the presence of discrete RNA isoforms with regions of overlap, while also allow mapping of TSS and CPAS. Finally, we performed Illumina Cap Analysis Gene Expression Sequencing (CAGE-Seq) (Murata et al., 2014) to map TSS by an orthologous approach (Figure 1A).

**Figure 1.**
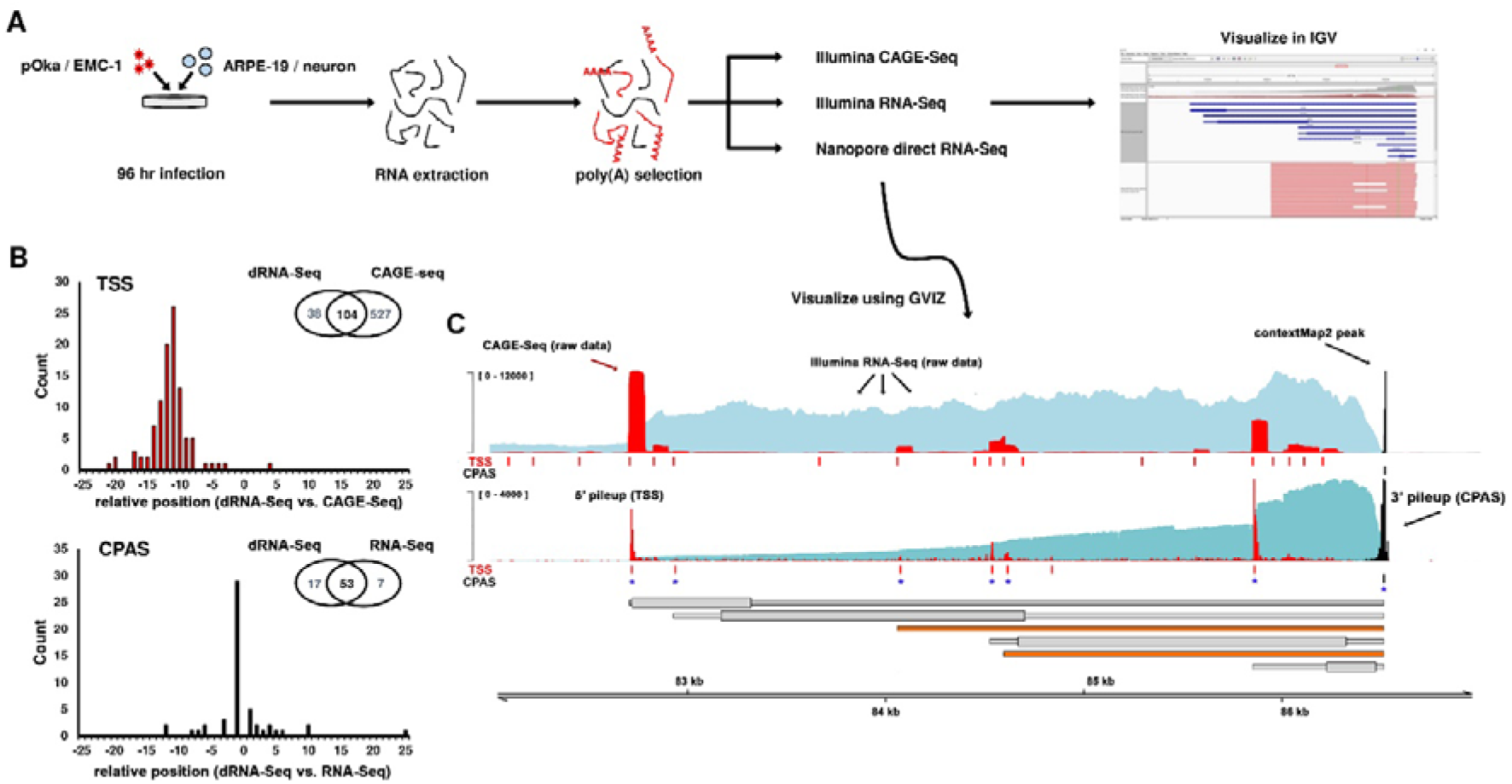
Decoding the complexity of the lytic VZV transcriptome. **(A)** Experimental strategy: ARPE-19 cells and hESC-derived neurons were infected with VZV EMC-1 (Clade 1) or -pOka (Clade 2) for 96 hrs. Total RNA was extracted and poly(A) fraction isolated for sequencing by Illumina CAGE-Seq, Illumina RNA-Seq and/or Nanopore dRNA-Seq. Sequence data were aligned against the VZV strain Dumas reference genome (Genbank Accession: NC_001348.1) and visualized using the Integrative Genomics Viewer (IGV) (Thorvaldsdóttir et al., 2013) and GVIZ (Hahne and Ivanek, 2016). **(B)** Transcription start sites (TSS) as well as cleavage and polyadenylation sites (CPAS) were identified in nanopore and Illumina datasets. Histograms show the distances observed between nanopore and Illumina predictions, while inset Venn diagrams indicate the numbers of sites identified and their conservation between datasets. **(C)** Integration of Illumina RNA-Seq, Illumina CAGE-Seq and Nanopore dRNA-Seq datasets. Coverage plots for Illumina RNA-Seq (light-blue), CAGE-Seq (red) and Nanopore dRNA-Seq (teal) are integrated with pileup data that maps TSS (red) and CPAS (black). Rows denoted by TSS and CPAS indicate positions of TSS and CPAS identified using HOMER software (Heinz et al., 2010) for nanopore dRNA-Seq and Illumina CAGE-Seq data and ContextMap2 software (Bonfert et al., 2015) for Illumina RNA-Seq data. Conserved TSS and CPAS are indicated with blue asterisks. RNA structures (grey) are inferred from these conserved sites. Wide and thin boxes indicate canonical coding sequence (CDS) domains and untranslated regions (UTRs), respectively. Novel identified RNAs (orange) are shown without predicted CDS domains.

TSS and CPAS estimates provided by dRNA-Seq data closely overlapped with those derived from our Illumina approaches (Figure 1B). TSS sensitivity was nearly four-fold higher in CAGE-Seq datasets compared to dRNA-Seq, with effectively all TSS uniquely found by CAGE-Seq being low abundance – likely reflecting artefacts derived from RNA processing (i.e. recapping of cleaved RNA) or a generalized dysregulation of transcription initiation accompanying the late stages of a viral infection. A total of 104 TSS overlapped between the dRNA-Seq and CAGE-Seq datasets, all of which were the most abundant TSS in both datasets (Table S1). Importantly, dRNA-Seq TSS estimates were located up to 20 nucleotides (nt) downstream (median 11 nt) of TSS derived via CAGE-Seq (Figure 1B). This difference is best explained by the presence of low-quality ends of dRNA-Seq reads that are not aligned when using local alignment strategies. CPAS sensitivity was higher in dRNA-Seq than RNA-Seq datasets (70 vs. 60 sites) with 53 sites detected by both approaches (Table S2). CPAS estimates provided by dRNA-Seq aligned closely with those derived from RNA-Seq data (median 1 nt difference, Figure 1B) due the 3’ → 5’ direction of dRNA-Seq. Finally, we reconstructed the VZV transcriptome using TSS and CPAS to define transcript structures followed by visual confirmation of read data to identify splice sites, define alternatively spliced transcripts, and examine read-through transcription (Figure 1C).

### Reannotation of the VZV transcriptome reveals alternative transcript isoforms and putative non-coding RNAs

In VZV pOka-infected ARPE-19 cells, we identified 136 distinct VZV RNAs that were readily detectable at 96 hpi. Along with defining the UTRs of 96 RNAs encoding the 71 canonical VZV ORFs, we also identified 40 additional RNAs (Figure 2, Table S3). To reduce confusion, we numbered all RNAs according to the respective canonical ORFs and delineated transcript isoforms encoding at least part of the same ORF by number. For instance, three transcripts are transcribed from the ORF0 locus and these are here referred to as VZV RNA 0-1, 0-2, and 0-3. The identified RNAs included transcripts encoding 5’ extended ORFs, 5’ truncated ORFs, 3’ extended ORFs, 3’ truncated ORFs, internally spliced variants, and putative non-coding RNAs (ncRNAs). Importantly, our study confirmed several previously described RNA isoforms such as a genome-termini spanning RNA that encodes ORF0 (RNA 0-3) (Kemble et al., 2000) and identified an additional 5’ truncated ORF0 RNA of unknown function (RNA 0-2) (Figure 3A). Similarly, extensive low-level internal splicing of ORF50 has previously been reported (Sadaoka et al., 2010) (RNAs 50-1 to 50-5) and was similarly observed here, supplemented by two additional 5’ truncated isoforms (RNAs 50-6 and 50-7) expressed at relatively high abundance (Figure 3B). Examples of previously undocumented RNAs include two new spliced ORF9 transcript isoforms and two internally spliced transcript isoforms encoding N-terminal ORF24 and ORF48 coding sequences (CDS) domains that are spliced into novel C-terminal domains (Figure 2). Finally, we confirmed expression of transcripts encoding the novel ORF9 and ORF48 variants in VZV-infected ARPE-19 cells by RT-PCR (Figure S1A-B).

**Figure 2.**
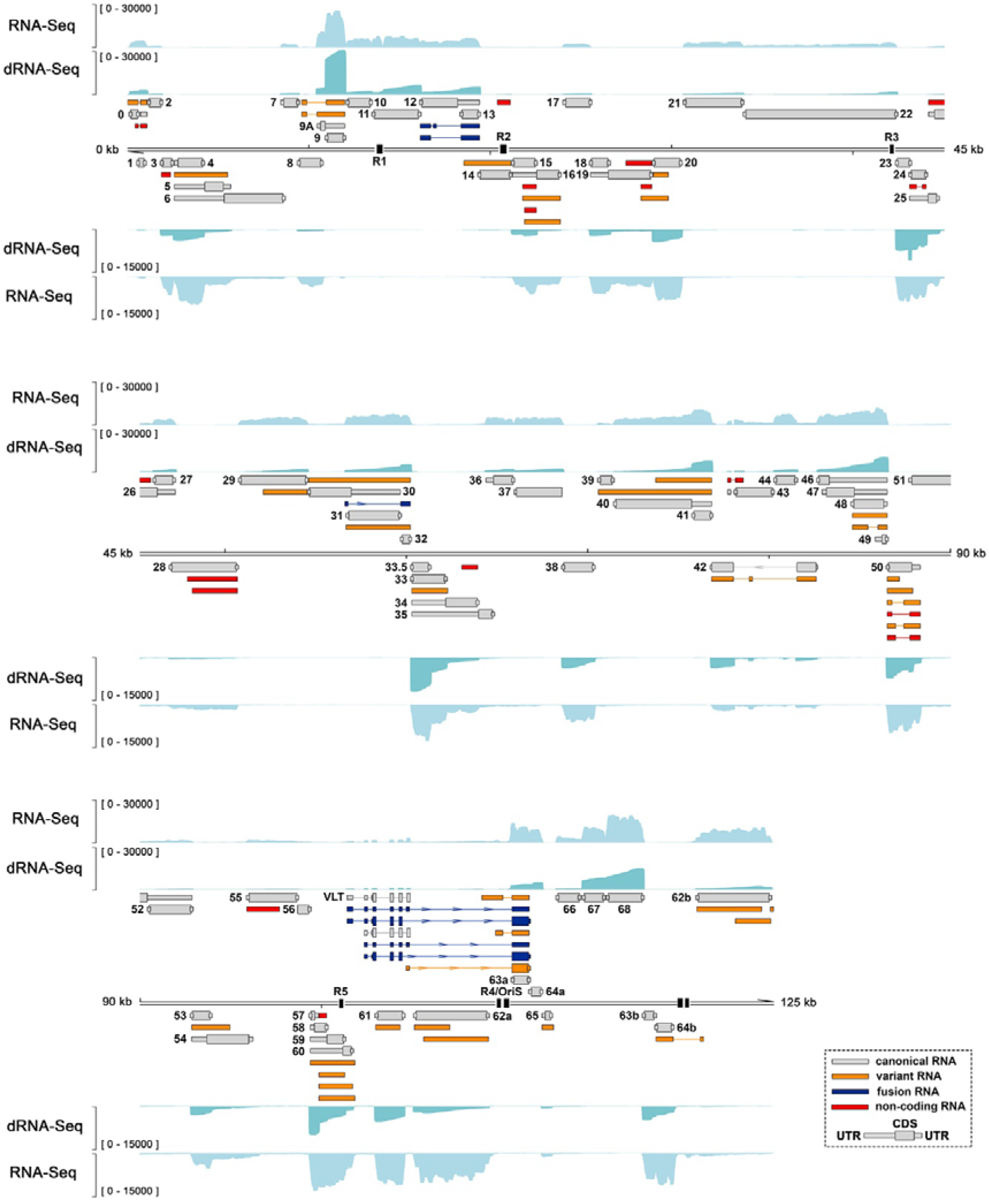
Reannotation of the lytic VZV transcriptome reveals novel viral transcript isoforms, fusion transcripts and putative noncoding RNAs. The reannotated lytic VZV transcriptome includes 71 transcripts encoding canonical ORFs with their UTRs defined (grey), 39 alternative isoforms of existing RNAs (orange), 7 fusion RNAs (dark blue) and 19 novel polyadenylated RNAs (red). Wide and thin boxes indicate canonical CDS domains and while thin boxes indicate UTRs, respectively. Absence of CDS regions indicates that the respective VZV RNA has an uncertain coding potential. Illumina RNA-Seq (light blue) and nanopore dRNA-Seq (teal) coverage plots are derived from ARPE-19 cells lytically infected with VZV strain pOka for 96 hr. Y-axis values indicate the maximum read depth of that track. See also Figures S1 and S2. Reiterative repeat regions R1-R5 and both copies of the OriS are shown as black boxes embedded in the genome track.

**Figure 3.**
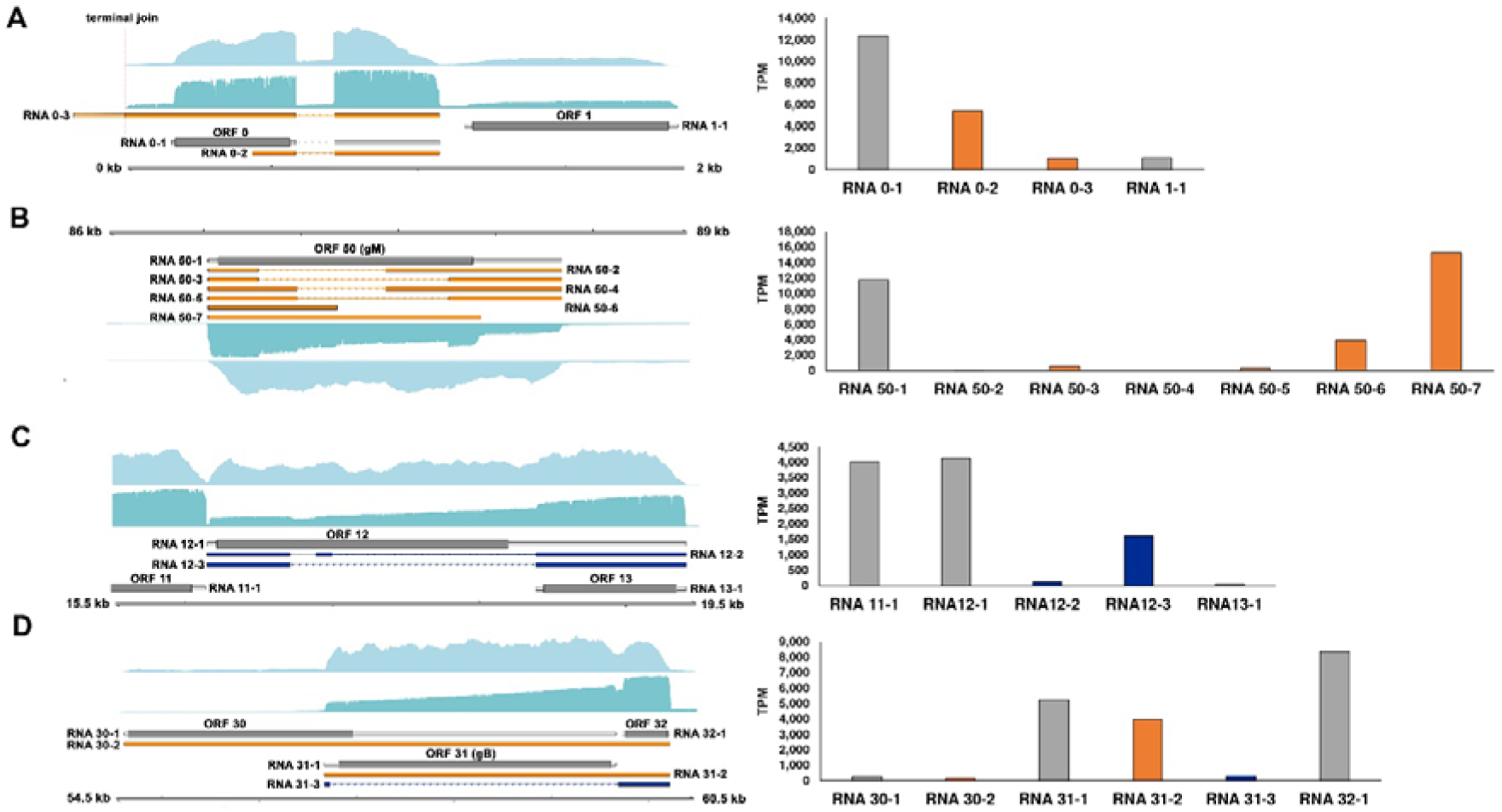
Examples of specific variant and fusion viral RNAs expressed during lytic VZV infection. **(A-D)** Examples of variant transcripts encoding ORF0 **(A)** and ORF50 **(B)** and novel fusion transcripts encoding (parts of) ORF12 and ORF13 **(C)**, and ORF31 and ORF32 **(D)**. Illumina RNA-Seq (light blue) and nanopore dRNA-Seq (teal) coverage plots are derived from ARPE-19 cells lytically infected with VZV strain pOka for 96 hr. Fusion RNAs are shown in dark blue, with variant RNAs shown in orange and canonical RNAs in grey. Hashed red bar indicates the position of the genome termini. Genome coordinates are shown while strand is indicated by placement of the genome track (below: top strand, above: bottom strand). Y-axes denote transcript per million (TPM) counts. See also Figure S4.

Fusion transcripts combine sequences from two or more distinct canonical viral transcripts, and mostly likely result from transcription termination occurring at an alternative CPAS downstream of the canonical CPAS, followed by internal splicing (Depledge et al., 2019). The resulting fusion transcripts are predicted to encode new proteins that contain fused domains from two or more distinct protein products. We identified seven VZV fusion transcripts in total. Four of these contain distinct fusion of transcripts encoding VLT and ORF63. Internal splicing of RNA 12-2 and RNA 12-3 transcripts yields two distinct splice variants that fuse parts of ORF12 and ORF13 CDS domains (Figure 3C). Additionally, we observed transcripts that fused the 5’ UTR of ORF31 transcripts to ORF32 transcripts, resulting in a transcript encoding pORF32 with an alternative 5’ UTR. We confirmed expression of transcripts encoding the ORF12-ORF13 and ORF31-ORF32 fusions in VZV-infected ARPE-19 cells by RT-PCR (Figure S1C-D).

Additionally, we discovered two novel polyadenylated VZV transcripts: RNA 13.5-1 and RNA 43-2. RNA 13.5-1 is 634 nt long, encodes two putative CDS domains (88 and 55 amino acids; aa), and is positioned antisense to the RNA 14-1 (encoding ORF14). Like RNA 14-1, RNA 13.5-1 stretches across the R2 reiterative region, a short repeat region that exhibits length variations between viral strains and within viral populations and thus leads to length variations in the encoded transcripts (Jensen et al., 2020) (Figures 2 and S1E). We also identified a highly expressed 3’ truncated RNA (RNA 43-2) that overlaps with the 5’ end of RNA 43-1 (encoding ORF43, Figures 2 and S1F). RNA 43-2 is a spliced 590 nt transcript that encodes only a short CDS domain (21 aa). Expression of both RNA 13.5-1 and RNA 43-2 in VZV-infected ARPE-19 cells was confirmed by RT-PCR (Figures S1E-F)

Finally, we used an *in-silico* approach to predict the coding potential of all 136 polyadenylated VZV RNAs (Table S3). The Coding Potential Calculator version 2 algorithm (CPC 2.0) (Kang et al., 2017) calculates the coding probability of a transcript based on its length, isoelectric point, and Fickett score of the longest CDS encoded. Of the 96 VZV RNAs encoding the 71 canonical ORFs, 89 were assigned a coding probability exceeding 90% with only two canonical RNAs – encoding the two smallest VZV proteins: pORF49 (81 aa) (Sadaoka et al., 2007) and pORF57 (71 aa) (Cox et al., 1998) – incorrectly predicted to be a non-coding transcript (Table S3). Of the 40 VZV RNAs encoding non-canonical products, 17 were predicted to be non-coding (Table S3), including two novel transcripts: RNA 13.5-1 (7%) and RNA 43-2 (13%) (Figure S2).

### The lytic VZV transcriptome is not influenced by viral strain or cell type

VZV genome sequences are highly conserved (Norberg et al., 2015), suggesting that strain-specific differences in coding capacity are likely minimal. To test this hypothesis, we infected ARPE-19 cells with VZV EMC-1 for 96 hrs and sequenced the poly(A) fraction of RNA by dRNA-Seq (Table S4). This enabled a comparative analysis of datasets obtained from ARPE-19 cells lytically infected with either VZV pOka or EMC-1 to determine whether either strain encodes unique transcripts (Figure S3). No such transcripts were identified at the RNA level, although we note that nucleotide level changes may still impact encoded proteins – as is exemplified by the N-terminal extended pORF0 uniquely present in pOka (Figure S4).

As VZV is capable of infecting diverse cell types including epithelial cells and neurons, we also determined if the VZV transcriptome remains similar between VZV pOka-infected ARPE-19 cells and human embryonic stem cell (hESC)-derived neurons (Sadaoka et al., 2016, 2017). Again, no cell-type specific novel VZV RNAs or variant VZV RNAs were identified (Figure S5), indicating that observed differences in VZV RNA expression levels and infectivity in distinct cell types (Baird et al., 2014; Sadaoka et al., 2017) is not due to the presence of cell-type specific VZV RNAs, but is likely driven by host cell factors.

### Overexpression of RNA 43-2 does not impair VZV replication in epithelial cells

Based on the high relative expression of RNA 43-2, its low protein coding potential and genomic location we hypothesized that it may function as a ncRNA involved in regulating the expression of the longer RNA 43-1 (encoding pORF43). ORF43 is an essential gene (Zhang et al., 2010), which putatively encodes for the capsid vertex component 1 and is postulated to be important for viral DNA encapsidation from the analogy to HSV UL17 (Toropova et al., 2011). To test this hypothesis, we generated three stable ARPE-19 cell lines expressing RNA 43-2, or an empty vector as control, and analyzed VZV EMC-1 replication at 48 and 72 hpi by flow cytometry and plaque assay. We did not observe any significant differences in number of VZV-infected cells, number of plaques nor plaque sizes between RNA 43-2 expressing cells and vehicle control cells (Figures S2A-B). Additionally, RNA 43-2 expression did not significantly reduce expression of RNA 43-1 in VZV-infected cells (*p* = 0.08 by Student’s t-test) (Figure S2C).

### Decoding the kinetic class of VZV transcripts in lytically infected cells

To determine the kinetic class of each lytic VZV RNA, cell-cycle synchronized ARPE-19 cells were infected with cell-free VZV EMC-1 and cultured for 12 or 24 hrs in the presence or absence of actinomycin D (ActD, transcription inhibitor), cycloheximide (CHX, translation inhibitor), or phosphonoacetic acid (PAA, inhibitor of the viral DNA polymerase), and viral RNAs were subsequently profiled using dRNA-seq and RNA-seq (Figure 4 & S6). Transcription of *IE* RNAs is not dependent on *de novo* viral protein production, whereas transcription of *E* RNAs depends on *IE* proteins. *L* RNAs are further subclassified into two kinetic classes, *leaky-late* (*LL*) and *true-late* (*TL*); *LL* RNAs are expressed at very low levels before, and *TL* RNAs exclusively after, viral DNA replication has commenced.

**Figure 4.**
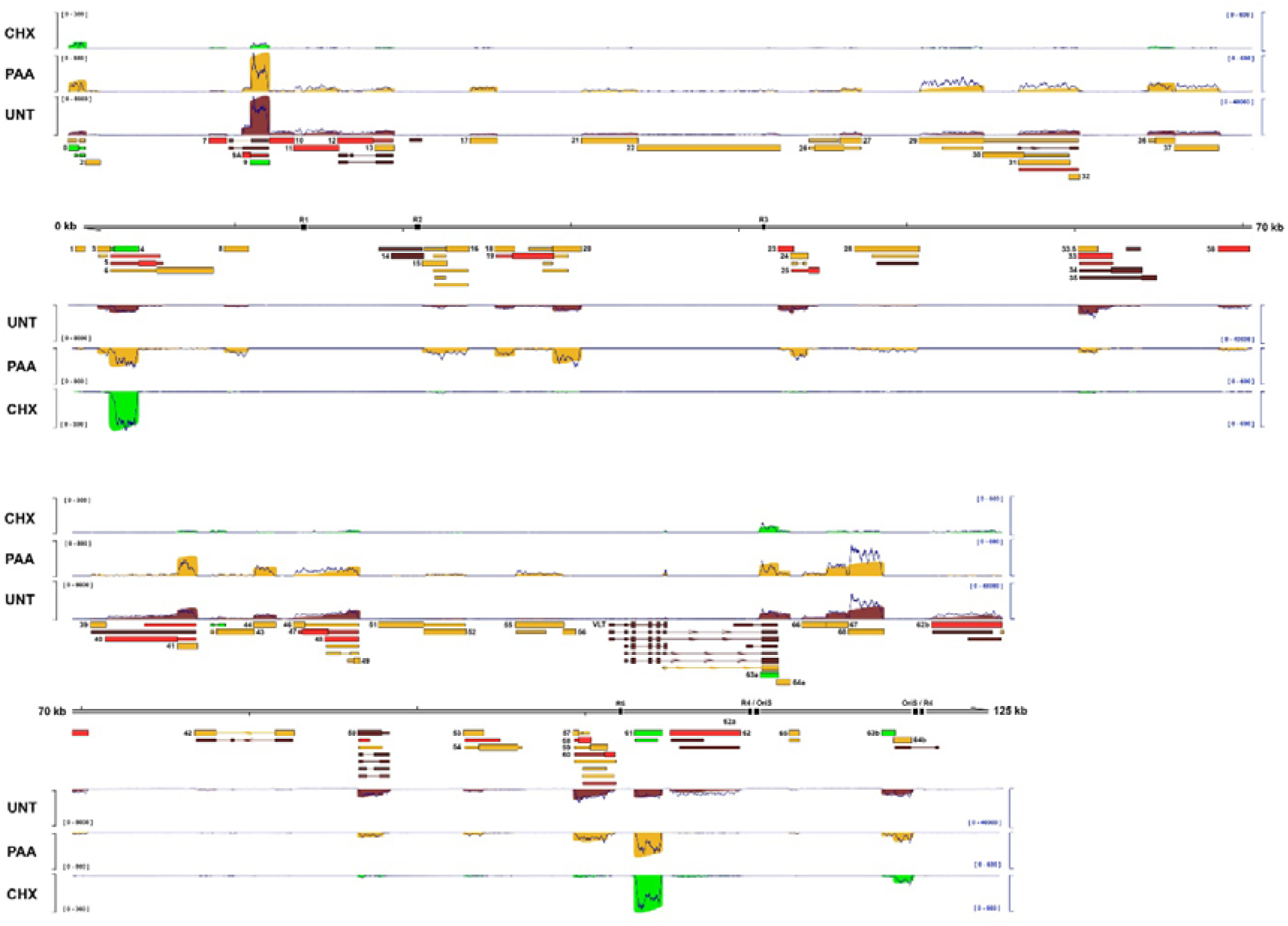
Decoding the kinetic class of VZV transcripts during lytic infection. Coverage plots derived from dRNA-Seq of ARPE-19 cells infected with cell-free VZV EMC-1 and treated with cycloheximide (CHX; green) for 12 hrs, phosphonoacetic acid (PAA; gold) for 24 hrs, or untreated (UNT; red) for 24 hrs. Corresponding Illumina RNA-Seq coverage plot for each condition are shown as dark blue line plots. Y-axis denotes coverage range (left side: nanopore dRNA-Seq, right-side: Illumina RNA-Seq). Reannotated VZV genome is shown in the middle track. CDS regions and UTRs are shown as wide and thin boxes, respectively. Reiterative repeat regions R1-R5 and both copies of the OriS are shown as black boxes embedded in the genome track. See also Figure S6. Canonical (grey), alternative (orange), fusion (dark blue) transcripts and putative ncRNAs (red) are shown with canonical CDS regions UTRs. Absence of CDS regions indicate that the respective RNAs have uncertain coding potential.

Given the very high sensitivity of our Illumina RNA-seq and Nanopore dRNA-seq analyses many VZV transcripts were detected in all experimental conditions except ActD-treated cells, albeit at vastly different abundancies (Table S3). Therefore, we first established objective criteria to classify VZV transcripts into distinct kinetic classes, based on their susceptibility to CHX treatment (to identify *IE* genes) and PAA treatment (to identify *E* and *L* genes). Taking advantage of the fact that one dRNA-Seq equals one RNA, we counted the number of reads mapping unambiguously to each of the 136 VZV RNAs for each sample. We subsequently calculated the relative expression level for each transcript by expressing each count as a fraction of the total VZV transcripts counts for that sample (Figure 5). Reads that could not be unambiguously assigned were excluded from this analysis. We defined *IE* VZV transcripts as those which accounted for an equal to or higher fraction of transcripts in the CHX treated sample than in the PAA or untreated samples (Figure 5). *E* RNAs were assigned as those which had a proportional distribution in PAA-treated samples of at least 50% of the untreated sample. *LL* transcripts were those which had a proportional distribution in PAA-treated samples of 5% – 50% of the untreated sample. *TL* transcripts were those with a proportional distribution in PAA-treated samples of less than 5% of the untreated sample and were effectively only detected in untreated samples.

**Figure 5.**
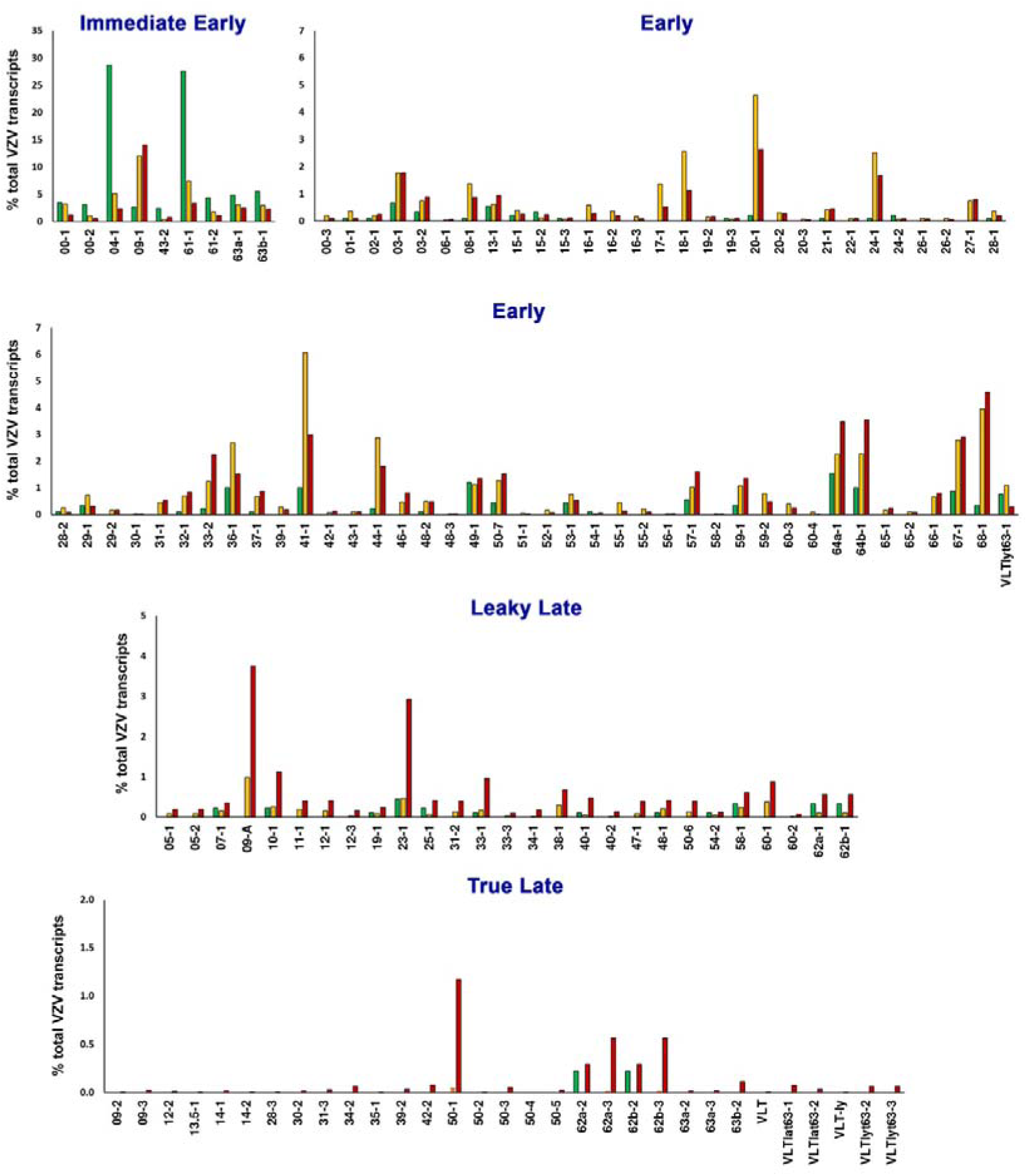
Transcriptional classification of VZV transcripts. The kinetic class of all viral RNAs included in our reannotated VZV genome was based on their differential expression in lytically VZV-infected ARPE-19 cells treated with cycloheximide (CHX; green), phosphonoacetic acid (PAA; gold), or untreated (UNT; red) (Figure 4). Viral *immediate-early* (*IE*) RNAs were assigned as those expressed proportionally highest in CHX-treated VZV-infected ARPE-19 cells. Viral *early* (*E*) RNAs were assigned as those expressed with proportional distribution in PAA-treated samples of at least 50% of the untreated VZV-infected ARPE-19 cells. Viral *leaky late* (*LL*) RNAs were those expressed at a proportional distribution in PAA-treated samples of at least 5% of the untreated VZV-infected ARPE-19 cells. Viral *true late* (*TL*) RNAs were those that were only detected in untreated VZV-infected ARPE-19 cells. Y-axis indicates the fractional representation of each specific RNA as percentage of the total number VZV transcripts detected in the respective (un)treated VZV-infected ARPE-19 cells.

Collectively, our approach showed that two VZV transcripts (RNA 4-1 and RNA 61-1, encoding pORF4 and pORF61, respectively) were expressed at very high levels (accounting for 60% of all VZV transcripts) in the absence of *de novo* protein production and, in agreement with prior studies (Moriuchi et al., 1993, 1994), were classified as *IE* transcripts (Figure 5). Six additional transcripts were expressed to high levels in CHX-treated samples relative to other conditions and were also assigned *IE* status. These included RNA 63-1 (encoding pORF63), RNA 0-1 (pORF0), RNA 0-2 (putative ncRNA), RNA 61-2 (N’ terminal truncated pORF61), and RNA 43-2 (putative ncRNA). We also observed that RNA 9-1 (pORF9) is expressed at similar levels as the transcripts above and provisionally classified it as *IE* but note that relative expression levels of this transcript increases throughout infection. pORF4, pORF61, and pORF63 are known transcriptional activators of VZV and canonical *IE* transcripts (Kost et al., 1995; Moriuchi et al., 1993, 1994), whereas the kinetic class of ORF0 has not been fully resolved (Koshizuka et al., 2010). Low level transcription of other VZV transcripts was observed but attributed to low-level transactivation by viral tegument proteins delivered from incoming virions. We also sequenced samples treated with ActD to control for the potential presence of residual background transcripts in the virus preparations and confirmed only minimal amounts of VZV transcripts to be present (Figure S6).

In total, 69 transcripts were classified as viral DNA replication insensitive *E* RNAs, including the experimentally validated transcripts encoding pORF28 and pORF29 (Yang et al., 2004). A further 27 transcripts were classed as *LL* and 31 transcripts as *TL*, the latter including both RNA 14-1 (pORF14) and VLT (pVLT), both of which have been experimentally confirmed previously (Depledge et al., 2018b; Storlie et al., 2006). Notably, about half of the VZV RNAs originating from the same locus were of different transcriptional class (Figure S6). Typically, the shortest RNA isoforms were of earlier kinetic class, with alternative TSS and CPAS usage increasing transcript diversity by producing longer alternative RNA isoforms at later stages of infection, e.g. RNA 9-1, 9A-1, 9-2, and 9-3 (Figure S6B).

### ORF62 transcripts are expressed with *Late* kinetics during lytic VZV infection

VZV transcripts RNA 62-1 and RNA 62-2 encode for, respectively, the major viral transcriptional activator protein, pORF62, and a predicted N-terminal truncated pORF62 variant. Surprisingly, our data indicate that expression of the ORF62 encoding RNAs is both dependent on *de novo* (viral) protein synthesis and viral DNA replication, thereby classifying these RNA 62 transcripts as *L*. This contradicts the current classification of ORF62 as an *IE* gene, although we note this classification was obtained by analogy to the function of its HSV-1 orthologue infected cell polypeptide 4 (ICP4) (Felser et al., 1988). To confirm and substantiate our findings, we analyzed the impact of viral DNA replication on ORF62-encoding RNA and protein expression in multiple VZV-susceptible cell types at 24 hpi. RT-qPCR analysis showed that PAA treatment was associated with an approximately 10-fold decrease in expression of *IE* transcripts encoding ORF61 and ORF63, consistent with the absence of VZV DNA replication and spread in culture, and >10,000-fold decrease in expression of *TL*, viral DNA replication-dependent, RNA VLTly (Figure 6A). Notably, expression of RNAs encoding ORF62 was more severely affected (∼500-fold reduction) by PAA treatment than *IE* RNAs encoding ORF61 and ORF63. Similar results were obtained for VZV strain pOka infected epithelial ARPE-19 cells, hESC-derived neurons and lung fibroblast MRC-5 cells (Figure S7A to C).

**Figure 6.**
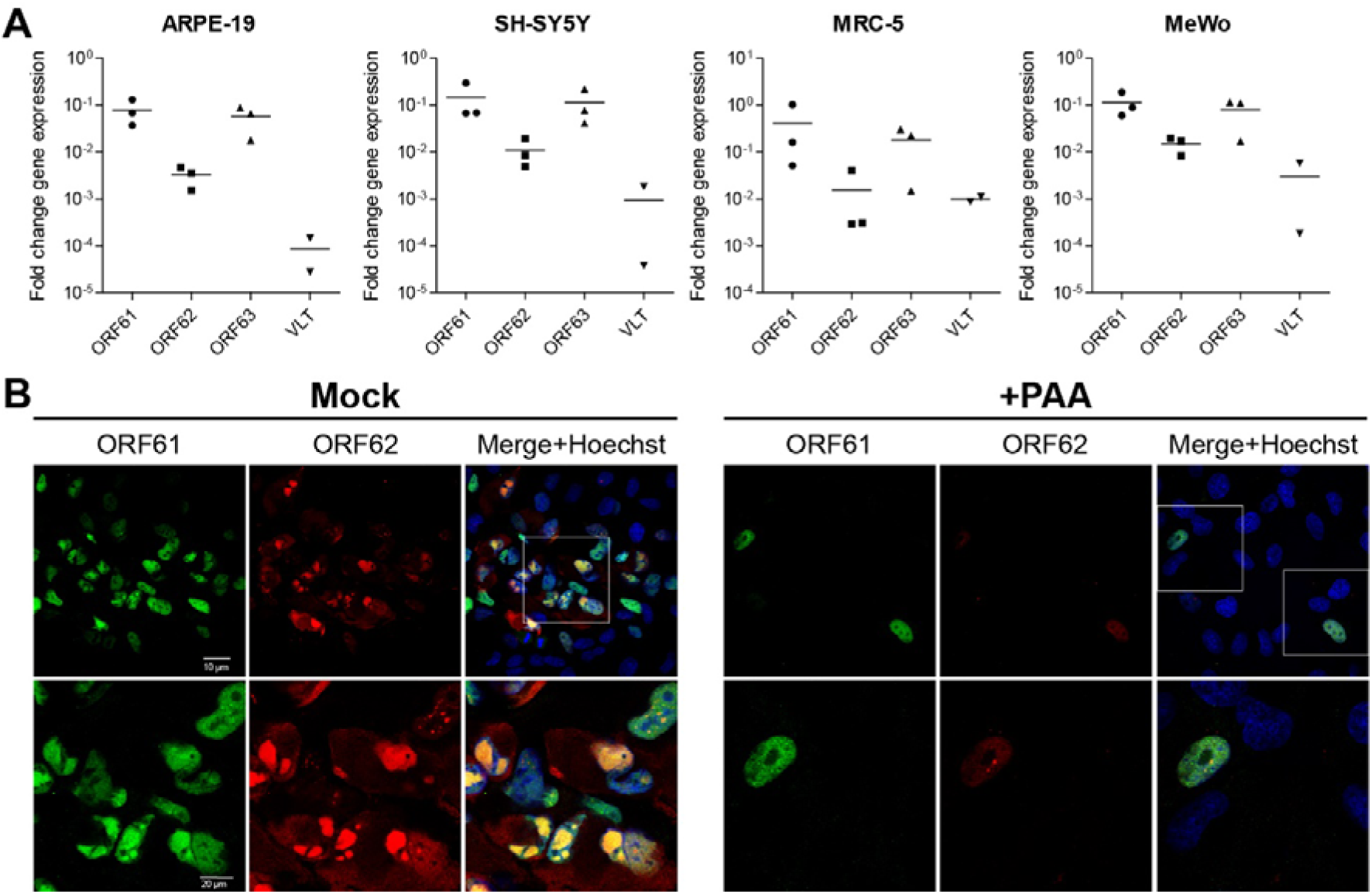
ORF62 transcripts are expressed with *Late* kinetics during lytic VZV infection. Cell-cycle synchronized human ARPE-19 (epithelial), SH-SY5Y (neuroblastoma), MRC-5 (fibroblast) and MeWo (melanoma) cells were infected with cell-free VZV EMC-1 for 24 hr in the presence or absence of PAA. **(A)** Fold change ORF61, ORF62 and ORF63 gene expression in PAA-treated samples relative to untreated samples (calibrator) and normalized to GAPDH using the 2^-ΔΔCt^ method. In 1 of 3 experiments, PAA-treatment reduced VLT expression to below detection limit. qPCR probes direct to ORF62 recognize all three transcript variants (RNA62-1, 62-2 and 62-3). **(B)** Representative images of lytically VZV-infected ARPE-19 cells stained by immunofluorescence for pORF61 (green) and pORF62 protein (red). Nuclei were counterstained with Hoechst 33342 (blue). Top row: 400x magnification, bottom rows: 1000x magnification. See also Figure S7.

Similarly, PAA treatment severely reduced the abundance and affected the cellular localization of pORF62 in VZV-infected ARPE-19 cells (Figure 6B). In the absence of PAA, both pORF61 and pORF62 were abundantly expressed in plaques of VZV-infected cells, with mostly diffuse nuclear pORF61 staining and pORF62 staining presenting as abundant globular nuclear and diffuse cytoplasmic staining (Figure 6B, left panels). Consistent with inhibition of VZV replication, no plaques were observed in PAA-treated cultures and infected cells were rare. The pORF61 staining pattern in infected cells was comparable between PAA-treated and untreated VZV-infected cells, whereas pORF62 staining intensity was severely reduced and showed weak, mostly diffuse nuclear staining with fewer intensely staining punctae (Figure 6B, right panels); possibly, reflecting incoming pORF62 originating from VZV virions. Identical IF staining results were observed in ARPE-19, MRC-5 and melanoma MeWo cells (Figure S7D and E, respectively). Overall, our data demonstrate that RNA 62-1, as well as RNA 62-2 and RNA 62-3, are expressed at low levels prior to viral DNA replication, with robust expression occurring only after viral DNA replication is initiated, consistent with the expression of *LL*, but not *IE* transcripts.

## Discussion

Understanding the full coding capacity of a given virus is crucial to understanding its biology. With the advent of new RNA-sequencing methodologies it has become clear that transcription of herpesvirus genomes is much more complex than previously anticipated. Here, we demonstrate that VZV is no exception and provide a comprehensive reannotation of the VZV transcriptome during lytic infection of human retinal pigment epithelial cells and hESC-derived neurons, incorporating data from two distinct VZV strains. By integrating RNA-Seq, CAGE-seq, dRNA-seq, we have resolved the architecture of the lytic VZV transcriptome. Specifically, we report the TSS and CPAS for all annotated VZV transcripts – including refinement of the 5’ UTRs and 3’ UTRs in RNAs encoding canonical ORFs – and show that VZV further diversifies its transcription by through the use of (1) additional TSS and CPAS, (2) disruption of transcription termination, and (3) alternative splicing. As a result, several transcript isoforms are expressed from the same locus and fusion RNAs are produced that modify UTRs and CDS of multiple transcripts. Given that 5’ UTR sequences influence the translational efficiency of the downstream CDS (Leppek et al., 2018), alternative UTR usage may provide the virus additional mechanism to regulate its protein expression throughout its infectious cycle. Collectively, this study defined 136 polyadenylated VZV RNAs that are expressed during lytic VZV infection, many of which are predicted to increase the diversity of the viral proteome.

Although the VZV genome is considered relatively stable, multiple strains currently co-circulate and recombine (Norberg et al., 2015; Tyler et al., 2007), potentially influencing the viral transcriptome. However, our comparison of the transcriptional landscape of a VZV clade 1 (strain EMC-1) and a clade 2 (strain pOka) virus revealed that no strain-specific lytic transcript isoforms exist in VZV-infected ARPE-19 cells. However, inter-strain differences in repeat variations could nevertheless impact transcription of RNA 11-1 (containing R1), RNA 13.5-1 (R2), RNA 14-1 (R2), RNA 22-1 (R3), RNA 63-2 (R4), RNA 63-3 (R4) and all RNA 59 and RNA 60 isoforms (R5) (Jensen et al., 2020). Similarly, strain-specific polymorphisms may function to extend coding domains such as the N-terminal extended pORF0 (RNA 0-3) in VZV pOka in comparison with other VZV clades (Figure S4). Additionally, while previous studies suggested that the VZV transcriptome is generally similar across diverse cell types, none had sufficient resolution or used the methodologies required to disentangle transcript structures (Baird et al., 2014; Depledge et al., 2018b; Jones et al., 2014). Here, we demonstrated that identical transcript isoforms were detected during lytic VZV infection of human retinal pigment epithelial cells and hESC-derived neurons. Thus, while VZV strain-specific polymorphisms and/or cell type may influence viral gene expression, their impact on the lytic VZV transcriptome structure appears to be small.

Many herpesviruses express ncRNAs during lytic and latent infections (Hancock and Skalsky, 2018). Previously, we identified the putative dual-function polyadenylated VZV RNA VLT, which encodes a protein expressed during lytic infection, but is also functional as an RNA, inhibiting ORF61 RNA expression in overexpression experiments (Depledge et al., 2018b). Here, we identified 17 additional polyadenylated VZV RNAs that are predicted to be non-coding. Diverse functions have been attributed to human ncRNAs, including the modification of antisense or overlapping transcription events (Pelechano and Steinmetz, 2013; Saxena and Carninci, 2011). The generation of functional VZV mutant viruses with impaired expression of identified putative ncRNAs is challenging due to overlap with other viral transcripts. Therefore, we have studied the function of VZV RNA 43-2 by means of RNA 43-2 overexpression followed by VZV superinfection. However, RNA 43-2 did not significantly reduce expression of the overlapping RNA 43-1 transcript nor influence VZV replication. These results most likely reflect the complexity of VZV transcript regulation during lytic infection, as putative ncRNA 43-2 is expressed earlier (*IE* kinetics) and at higher abundance than RNA 43-1 (*E* kinetics). Considering the multitude of ncRNAs expressed by other herpesviruses and their crucial roles during infection (Hancock and Skalsky, 2018), delineating the functional importance of VZV ncRNAs should be considered a research.

Twenty-eight of 136 VZV transcripts are (multiply) spliced. Strikingly, the majority of splicing events occur in a hypercomplex region of the VZV genome encoding both VLT and ORF61. We have previously shown this locus to be characterized by extensive alternative splicing (Depledge et al., 2018b) and here report the presence of multiple transcripts that variously encode a fusion protein of pVLT and pORF63 (pVLT-ORF63) or an N’ terminal extended pORF63 (pORF63-N+). The coding potential of pVLT-ORF63 or pORF63-N+ and functional consequences of (and requirement for) these fusion transcripts during reactivation from latency is described in a separate study (Sadaoka *et al. unpublished data*), while their functional role(s) during lytic infection are under investigation. Except for VLTlyt63-1, all spliced VZV RNAs used the canonical splice donor (GT) site (Figure S8A). The splice donor sites were highly enriched for C/A (−3 position), A (−2), G (−1) and A (+1). All spliced VZV RNAs used the canonical splice acceptor site (AG), often flanked by C (−1) and G/A (+1) (Figure S8B). Consistent with the use of the cellular splicing machinery to process viral pre-mRNAs, VZV consensus splice donor and acceptor sites closely resemble those of the human transcriptome (Zhang et al., 2007).

Herpesvirus transcripts are traditionally assigned kinetic classes based on their temporal expression pattern and dependence on *de novo* protein synthesis or viral DNA replication. Here, we provide a transcriptome-wide classification of VZV transcripts during lytic infection of epithelial cells and identified that multiple transcripts originating from the same locus could either share the same kinetic class (e.g. RNAs 15-1, 15-2, and 15-3) or be of different kinetic class (e.g. RNAs 9-1, 9-2, 9-3, and 9A-1) (Table S3 and Figure S6). While the biological impetus for this is not clear, it seems likely that TSS and CPAS usage are dynamically regulated during lytic infection. For example, RNA 43-1 (*E,* pORF43) and the putative ncRNA 43-2 (*IE*) diverge in CPAS usage and expression kinetics (Figure 5). Similarly, multiple transcripts encoding pORF63 utilize different TSS are expressed as different temporal classes and varying abundancies, with canonical *IE* RNA 63-1 being most abundant, followed by *E* RNA VLTlyt63-1 and low quantities of two *TL* RNAs 63-2 and 63-3 (Figure 5).

Finally, our data provide novel insight into the expression of *IE* transcripts and the role of pORF62 during lytic VZV infection. The most abundantly expressed VZV *IE* transcripts, produced in the absence of new protein synthesis, encode for pORF4 and pORF61, which are also the earliest proteins detected in during lytic VZV infection (Ouwendijk et al., 2020). Interestingly, prior studies of the RNA 4-1 and RNA 61-1 promoter regions have shown that their efficient transactivation is dependent on pORF62 (Michael et al., 1998; Wang et al., 2009), a major component of the viral tegument (Kinchington et al., 1992). However, our data indicate that abundant expression of VZV RNA 62-1 (pORF62), as well as RNAs 62-2 and 62-3 is dependent on viral DNA replication (*LL* and *TL* kinetics), suggesting that tegument-derived pORF62 but not *de novo* pORF62 transactivates RNA 4-1 and RNA 61-1 expression at least during establishment of lytic infection cycle. Classification of RNA 62-1 (pORF62) as *LL* is also supported by prior observations that a marked increase in pORF62 abundance occurs only after DNA replication had commenced (Reichelt et al., 2009). Importantly, our findings do not exclude any of the previously assigned functions of pORF62, most notably its function as a major transcriptional regulator of VZV transcripts (Perera et al., 1992; Ruyechan et al., 2003; Yang et al., 2006). However, future studies aimed at better understanding the regulation of VZV transcript expression, and the distinct roles of newly produced RNA 62 isoforms, pORF62, and tegument-derived pORF62 are warranted.

In summary, this study describes the detailed analysis of the VZV transcriptome architecture and kinetic classification of viral transcripts in the context of lytic VZV infection. We provide these data as a comprehensive resource that will facilitate functional studies of coding RNAs and their protein products, the role of non-coding RNAs, and the regulation of VZV transcription and translation during lytic infection.

## Acknowledgements

We extend special thanks to Ian Mohr (New York University School of Medicine) for support of D.P.D. in part through National Institutes of Health (NIH) grants R01-AI073898 and R01-GM056927. S.E.B. was in part supported by the Nederlandse Organisatie voor Wetenschappelijk Onderzoek (NWO) project 022.005.032. This work was supported in part by the Takeda Science Foundation, Daiichi Sankyo Foundation of Life Science, Japan Society for the Promotion of Science (JSPS KAKENHI JP17K008858, JP16H06429 and JP16K21723) and the Ministry of Education, Culture, Sports, Science and Technology (MEXT KAKENHI JP17H05816) (T.S.). J.B acknowledges support from the National Institute for Health Research University College London Hospitals Biomedical Research Centre. The computational requirements for this work were supported in part by the NYU Langone High Performance Computing (HPC) Core’s resources and personnel.

## Author Contributions

W.J.D.O. and D.P.D. conceived of the project; W.J.D.O., D.P.D., and T.S. designed the experiments with additional input from G.M.G.M.V.; S.E.B., T.S., W.J.D.O., and D.P.D. performed the experiments and analyzed the data; S.E.B., T.S., W.J.D.O., and D.P.D. wrote the manuscript; All authors read, edited, and approved the final paper.

## Methods

### Cells and viruses

Human retinal pigmented epithelium ARPE-19 cells [American Type Culture Collection (ATCC) CRL-2302] were grown in a 1:1 (v/v) mixture of DMEM (Lonza) and Ham’s F12 (Gibco) medium supplemented with 10% heat-inactivated fetal bovine serum (FBS; Lonza) and 0.6 mg/mL L-sodium glutamate (Lonza) or in DMEM/F-12+GlutaMAX-I (Thermo Fisher Scientific) supplemented with heat-inactivated 8% FBS (Sigma-Aldrich). Human neuroblastoma SH-SY5Y cells were grown in a 1:1 (v/v) mixture of EMEM with EBSS (Lonza) and Ham’s F12 (Gibco) medium supplemented with 15% FBS (Lonza), L-sodium glutamate, penicillin-streptomycin, non-essential amino acids (MP biomedicals) and natrium-bicarbonate (Lonza). Human embryonic lung fibroblasts MRC-5 and human melanoma MeWo cells were cultured in DMEM (Lonza), supplemented with 10% FBS, L-sodium glutamate and penicillin-streptomycin. Human embryonic stem cell (hESC; H9)-derived neural stem cells (NSC) (Thermo Fisher Scientific) were cultured, propagated and differentiated into neurons as described previously (Sadaoka et al., 2020). Cell cultures were maintained at 37°C in a humidified CO_2_ incubator. VZV strain pOka (parental Oka) was maintained in, and the cell-free virus was prepared from, ARPE-19 cells as described previously for MRC-5 cells (Sadaoka et al., 2007). VZV strain EMC-1 is a low-passage clinical isolate, was cultured on ARPE-19 cells and cell-free VZV was extracted as described (Lenac Rovis et al., 2013; Ouwendijk et al., 2014). Cell-free EMC-1 was freshly harvested on the day of use and pretreated with DNAse I, RNAse T1 and RNAse A (all: Thermo Fisher Scientific) for 30 min at 37°C prior to infection.

### RNA extraction and cDNA synthesis

ARPE-19 cells were infected with cell-associated VZV EMC-1 by co-cultivation of uninfected and VZV EMC-1 infected ARPE-19 cells at an 8:1 cell ratio for 96 hrs. Alternatively, ARPE-19, SH-SY5Y, MRC-5 and MeWo cells were infected with cell-free VZV EMC-1 for indicated time. Cells were harvested in 1 mL TRIzol (Thermo Fisher Scientific), mixed with 200 μL chloroform and centrifuged for 15 min at 12,000xg at 4°C. RNA was isolated from the aqueous phase using the RNeasy Mini kit (Qiagen) according to manufacturer’s instructions, including on-column DNase I treatment, as described (Ouwendijk et al., 2013). RNA concentration and integrity were analyzed using a Nanodrop spectrophotometer (Thermo Fisher Scientific), and RNA was subjected to a second round of DNAse treatment using the TURBO DNA-free kit (Ambion) according to manufacturer’s instructions. For cDNA synthesis maximum 5 µg RNA was reverse transcribed using Superscript IV reverse transcriptase and oligo(dT) primers (Thermo Fisher Scientific) (RT+). As control, the same reaction was performed without reverse transcriptase (RT-). Alternatively, RNA was isolated using the FavorPrep Blood/Cultured Cell Total RNA Mini Kit (Favorgen Biotech) in combination with the NucleoSpin RNA/DNA buffer set (Macherey-Nagel). DNA was first eluted from the column in 100 µL DNA elution buffer and subsequently the column was treated with recombinant DNase I (20 units/100 µL; Roche Diagnostics) for 30 min at 37°C and finally RNA was eluted in 50 µL nuclease free water. RNA was directly treated with Baseline-ZERO DNase (2.5 units/50 µL; Epicentre) for 30 min at 37°C. cDNA was synthesized with 12 µL of RNA and anchored oligo(dT)18 primer in a 20 µL reaction using the Transcriptor First Strand cDNA synthesis kit at 55°C for 30 min for reverse transcriptase reaction (Roche Diagnostics).

### PCR and sequence analysis

PCR was performed on RT+ and RT-cDNA reactions using Amplitaq Gold DNA Polymerase (Thermo Fisher Scientific) and primer pairs corresponding to each newly identified VZV transcript (Table S5). Primers were directed to the predicted 5’ and 3’ end of each transcript so that newly identified transcripts were completely amplified from 5’→3’ end. For ORF9 variants and ORF48 additional reverse primers were used to confirm splice junctions (Table S5). PCR amplification was performed as follows: initial denaturation at 95°C for 10 min, followed by 40 cycles of alternating denaturation (30 sec, 95°C), primer annealing (30 sec at the appropriate temperature; Table S5), and subsequently primer extension (1 min / 1,000bp, 72°C; Table S5). Final extension step of 10 min at 72°C was included. To amplify ORF13.5 each dNTP, including equimolar amounts of dGTP and 7-deaza-GTP (New England Biolabs), at a concentration of 200 µM was used (Maertzdorf et al., 1999). PCR amplification of ORF13.5 was performed as follows: initial touchup PCR from 58°C→70°C using a transcript specific forward primer and an anchored primer on the poly(A) tail. Subsequently, semi-nested PCR was performed using the same forward primer and a reverse primer within the transcript using standard PCR protocol. Amplicons were purified from gel using the QIAquick Gel Extraction Kit (Qiagen) and sequenced using the BigDye v3.1 Cycle Sequencing Kit (Applied Biosciences) with corresponding forward and reverse primer on the ABI Prism 3130 XL Genetic Analyser.

### Plasmid construction and generation of stable cell lines

The RNA 43-2 transcript sequence (77,775 - 78,619 excluding the intron at 77,869 - 78,149; strain Dumas, NC_001348.1) was amplified using cDNA from VZV EMC-1 infected ARPE-19 cells and primers NheI_ORF43-5_Fw and XhoI_ORF43.5_Rv (Table S5). Amplicon was digested with NheI and XhoI and cloned into pcDNA3.1. Three independent batches of ARPE-19 cells were transfected with either pcDNA3.1/empty or pcDNA3.1/RNA 43-2 using polyethylenimine (PEI). After 2 days, cells were incubated with 1 mg/mL geneticin and cultured for at least 3 weeks to select for transfected cells. Subsequently, DNA was isolated using the QiaAmp DNA Mini kit according to manufacturer’s instructions and presence of RNA 43-2 DNA was confirmed by PCR. Next, RNA was isolated as described above, and expression of RNA 43-2 was confirmed by RT-PCR.

### Flow cytometry

ARPE-19 cells stably transfected with pcDNA3.1/empty or pcDNA3.1/RNA 43-2 were plated one day prior to infection in a 48-well plate. Cells were infected with cell-free VZV EMC-1 (multiplicity of infection, MOI = 0.01), harvested at 48 hpi or 72 hpi, fixed and permeablized with BD Cytofix/Cytoperm, stained for VZV glycoprotein E (gE) (MAB8612, Millipore) in BD PermWash, labeled with secondary APC-conjugated goat anti-mouse Ig antibody (BD Biosciences). Frequency of VZV-infected (i.e. APC positive) cells was measured on a BD FACSLyric flow cytometer and analyzed using FlowJo software (BD Biosciences).

### Plaque assay

Confluent monolayers of stable pcDNA3.1-empty or pcDNA3.1-RNA 43-2 cells grown in a 12-wells plate were infected with 2,000 plaque forming units (PFU)/well VZV EMC-1. At 72 hrs post-infection plates were washed and fixed with 4% paraformaldehyde (PFA) in PBS. Subsequently plates were permeabilized using 0.1% Triton-X100 in PBS for 10 min, blocked with 5% normal goat serum in PBS-0.05% Tween-20 (PBS-T) for 30 min, incubated with mouse anti-VZV gE antibody (MAB8612) diluted in PBS-T containing 0.1% BSA for 1 hr at room temperature. Cells were washed with PBS-T, incubated for 1 hr with polyclonal rabbit-anti-mouse Ig antibody (Dako) in PBS-T + 0.1% BSA, washed and stained with Alexa Fluor 488 (AF488)-conjugated goat anti-rabbit Ig (H+L) antibody (Thermo-Fisher) in PBS-T. Plates were measured using the Immunospot S6 Ultimate UV Image Analyzer and plaque size was determined using Immunospot software (Cellular Technology Limited).

### Kinetic class of VZV genes

ARPE-19 cells were synchronized in the cell cycle using a double thymidine block approach (Ma and Poon, 2016). Briefly, ARPE-19 cells were seeded at semi-confluence in 12-wells plates and next day medium was replaced for growth medium containing 2mM thymidine (Sigma-Aldrich). After 24 hrs, medium was replaced for normal growth medium for 8 hrs, after which medium was changed back to thymidine containing medium. 30 min prior to infection cells were released from thymidine by replacing the medium with regular culture medium. Cells were infected with freshly harvested cell-free VZV EMC-1 using spin-inoculation for 15 min at 1,000xg (MOI after spin-inoculation = 0.1 - 0.2). Cells were incubated for 45 min at 37°C, after which the inoculum was replaced for fresh medium, medium with 10 µg/mL Actinomycin D (ActD; Sigma-ALdich), medium with 50 μg/mL cycloheximide (CHX; C4859, Sigma-Aldrich) or medium with 400 µg/mL phosphonoacetic acid (PAA; Sigma-Aldrich). Infected cells were harvested in 500 μL TRIzol at 12 hpi (ActD and CHX) or 24 hpi (untreated and PAA) for RNA extraction. Alternatively, ARPE-19 cells, MRC-5 cells (1 x 10^5^ cells/well on 24-well plate) and hESC-derived neurons (1 x 10^5^ cells/well on 24-well plate as NSC and differentiated for 18 days) were infected with cell-free VZV pOka (20 µL of 4 x 10^4^ PFU/mL) in the presence or absence of phosphonophormic acid (200 µg/mL) (Sigma-Aldrich) for 1 hr, the inoculum was replaced for fresh media with phosphonophormic acid (200 µg/mL) and cultures were maintained for 24 hrs.

### Quantitative PCR analysis

Quantitative Taqman real-time PCR (qPCR) was performed in duplicate on RT- and RT+ cDNA using 4x Taqman Fast Advanced Master mix (Applied Biosystems) on a 7500 Taqman PCR system. Primer-probe sets directed to ORFs 61, 62, 63 and VLT have been described previously (Depledge et al., 2018b; Ouwendijk et al., 2012) and those directed to ORF43 are described in Table S7. Alternatively, cDNAs were subjected to qPCR using KOD SYBR qPCR Mix (TOYOBO) in the StepOnePlus Real-time PCR system (Thermo Fisher Scientific) (1 µL of cDNA per 10 µL reaction). All primer sets used for SYBR Green chemistry (Table S6) were first confirmed for the amplification rate (98-100%) using 10-10^6^ copies (10-fold dilution) of pOka-BAC genome or VLT plasmid (Depledge et al., 2018b) and the lack of non-specific amplification using water. The qPCR program is as follows; 95°C for 2 min (1 cycle), 95°C for 10 sec and 60°C 15 sec (40 cycles), and 60 to 95°C for a dissociation curve analysis. Data is presented as relative VZV level to cellular beta-actin defined as 2^-(Ct-value VZV gene - Ct-value beta-actin)^.

### Immunofluorescence staining

ARPE-19 cells were plated on glass coverslips in 24-wells plates one day prior to infection. Cells were inoculated with freshly harvested cell-free VZV EMC-1 in medium with or without 400 µg/mL PAA and incubated for 24 hrs. Infected cells were fixed with 4% PFA, permeabilized for 10 min with 0.1% Triton-X100 in PBS, blocked with 5% goat serum diluted in 0.2% gelatin-PBS solution and incubated on 30 µL 0.2% gelatin-PBS droplets containing primary antibody overnight at 4°C. The following primary antibodies were used: anti-pORF61 antibody (1:1000, gift from Dr. P. Kinchington) and monoclonal mouse anti-pORF62 antibody (1:200) (Lenac Rovis et al., 2013). Cells were washed 3-times with 0.2% gelatin-PBS and incubated for 1hr at room temperature with secondary antibodies diluted in 0.2% gelatin-PBS. The following secondary antibodies were used: AF488-conjugated goat anti-rabbit Ig (H+L) antibody (1:500; Thermo-Fisher) and AF594-conjugated goat anti-mouse Ig (H+L) antibody (1:500; Thermo-Fisher). Cells were washed once with 0.2% gelatin-PBS, washed once with PBS, incubated with a 1:1000 dilution of Hoechst 33342 (Life Technologies, 20 mM) in PBS for 5 min, washed with PBS and mounted using Prolong Gold Antifade Mounting medium (Thermo Fisher). Stained cells were analyzed using a Zeiss LSM 700 confocal laser scanning microscope (Zeiss) with a magnification of 400x or 1,000x. Photoshop CC 2019 software (Adobe) was used to adjust brightness and contrast.

### Illumina RNA sequencing and analysis

Stranded RNA libraries were prepared from poly(A)-selected RNA using the NEBNext® Ultra™ II Directional RNA Library Prep Kit for Illumina (New England Biolabs) and sequenced using a NextSeq 550. Sequence reads were trimmed using TrimGalore (https://www.bioinformatics.babraham.ac.uk/projects/trim_galore/) (--paired --length 30 –quality 30) and aligned against the VZV reference genome (strain Dumas, NC_001348.1) using BBMAP (https://sourceforge.net/projects/bbmap/) with post-alignment processing performed using SAMtools (Li et al., 2009) and BEDtools (Quinlan and Hall, 2010) to generate BEDGRAPH and BED12 files. Candidate CPAS were identified using ContextMap2 (Bonfert et al., 2015) (-aligner_name bowtie --polyA –strand-specific) with sequence reads aligned to VZV Dumas under default parameters by BWA (Li and Durbin, 2009).

### Cap analysis of gene expression (CAGE) sequencing and analysis

Two biological replicates of total RNA were extracted from ARPE-19 cells infected with VZV pOka for 96 hrs and CAGE-Seq libraries prepared by DNAFORM (Yokohama, Japan) and subsequently sequenced using an Illumina NextSeq 550, as previously described (Murata et al., 2014). Resulting sequence reads were trimmed (--length 30 --q 30 --clip_R1 1) using TrimGalore prior to alignment against the VZV Dumas genome using BBMAP. Post-alignment processing was performed using SAMtools and BEDtools. TSS were identified using the HOMER findPeaks module (-style tss -localSize 100 -size 15). Only TSS present in both biological replicates were retained for analysis.

### Nanopore direct RNA sequencing

For each biological sample, up to 1,000 ng of poly(A) RNA was isolated from up to 50 µg of total RNA using the Dynabeads™ mRNA Purification Kit (Invitrogen, 61006). Isolated poly(A) RNA was subsequently spiked with 0.3 µL of a synthetic Enolase 2 (ENO2) calibration RNA (Oxford Nanopore Technologies Ltd.) and dRNA-Seq libraries prepared as described previously (Depledge et al., 2019). Sequencing was performed on a MinION MkIb using R9.4.1 (rev D) flow cells (Oxford Nanopore Technologies Ltd.) for 18 – 44 hrs (one library per flowcell) and yielded between 720,000 – 1,290,000 reads per dataset (Table S4). Raw fast5 datasets were then basecalled using Guppy v3.2.2 (-f FLO-MIN106 -k SQK-RNA002) with only reads in the pass folder used for subsequent analyses. Sequence reads were aligned against the VZV Dumas genome, using MiniMap2 (Li, 2018) (-ax splice -k14 -uf --secondary=no), with subsequent parsing through SAMtools and BEDtools. Here sequence reads were filtered to retain only primary alignments (Alignment flag 0 (top strand) or 16 (bottom strand)).

### Splice junction correction in dRNA-Seq alignments

Illumina-assisted correction of splice junctions in RNA-Seq data was performed using FLAIR v1.3 (Tang et al., 2020) in a stranded manner. Briefly, Illumina reads aligning to the VZV Dumas genome were split according to orientation and mapping strand [-f83 & -f163 (forward) and -f99 & -f147 (reverse) ] and used to produce strand-specific junction files that were filtered to remove junctions supported by less than 50 Illumina reads. Direct RNA-Seq reads were similarly aligned to the Ad5 genome and separated according to orientation [-F4095 (forward) and - f16 (reverse)] prior to correction using the FLAIR correct module (default parameters). FLAIR-corrected alignments were used for all subsequent downstream analyses.

### TSS and CPAS identification in dRNA-Seq data

TSS and CPAS were identified by parsing SAM files to BED12 files in a strand-specific manner using BEDtools, and then truncating each aligned sequence read to its 5’ or 3’ termini for TSS and CPAS identification, respectively. Peak regions containing TSS and CPAS were identified using the HOMER findpeaks module (-o auto -style tss) using a --localSize of 100 and 500 and --size of 15 and 50 for TSS and CPAS, respectively. TSS peaks were compared against Illumina annotated splice sites to identify and remove peak artefacts derived from local alignment errors around splice junctions. 38 putative TSS identified in the dRNA-Seq dataset alone were flagged as artefacts and removed. Each of these TSS mapped precisely to a splice acceptor within a spliced RNA and closer inspection of the reads showed these to result from local alignment processes (Depledge et al., 2019).

### Generating RNA abundance counts from dRNA-Seq data

Using the updated VZV strain Dumas annotation presented here, we generated a transcriptome database by parsing our GFF3 file to a BED12 file using the gff3ToGenePred and genePredtoBED functions within UCSCutils (https://github.com/itsvenu/UCSC-Utils-Download) and subsequently extracting a fasta sequence for each annotated RNA using the getfasta function within BEDtools. dRNA-Seq reads were then aligned against the transcriptome database using parameters optimized for transcriptome-level alignment (minimap2 -ax map-ont -p 0.99). RNA counts were generated by counting alignments against a given RNA only if the alignment 5’ end was located within the first 50 nt of the RNA and the alignment was not marked as supplementary.

### *In silico* prediction of coding potential

CPC 2.0 (Kang et al., 2017) was used to examine the coding potential of all VZV RNAs defined in this study (Table S3). Note that RNAs were excluded from CPC 2.0 analysis and defined at putatively non-coding if no proteins greater than fifty amino acids in length were encoded.

### Data visualization

Figures associated with this study were generated using the R packages Gviz (Hahne and Ivanek, 2016) and GenomicRanges (Lawrence et al., 2013).

### Data availability

All sequencing datasets associated with this study are available via the European Nucleotide Archive under the accession PRJEB36978. Analysed datasets generated as part of this study, including a database of transcripts, BED12 alignment files, and GFF3 files describing our VZV annotation are freely available at https://github.com/dandepledge/vzv-2.0.

## Supplemental Information titles and legends

**Figure S1. Related to Figure 2.**
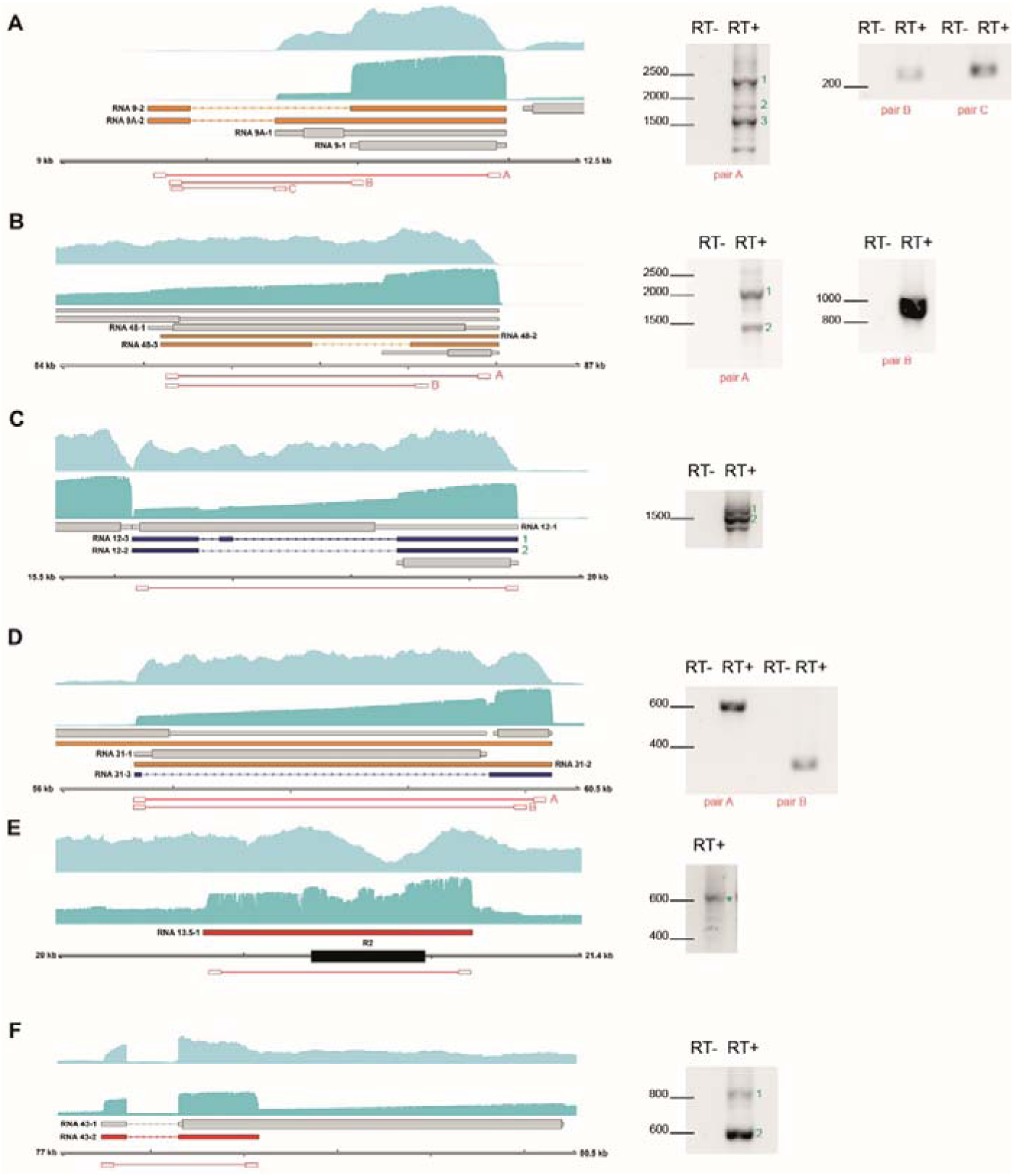
Confirmation of selected novel viral RNAs expressed during lytic VZV infection. The structure and location of selected VZV transcripts are shown alongside gel images confirming their expression. Illumina RNA-Seq (light blue) and Nanopore dRNA-Seq (teal) coverage plots are derived from 96 hr cell-associated infections of ARPE-19 epithelial cells. Transcript colours denote their status as canonical (grey), variant (orange), fusion (blue), or putatively non-coding (red). RT-PCR was performed on RNA extracted from ARPE-19 cells infected with cell-free VZV EMC-1 for 72 hours. RT+ / RT-: reverse transcriptase added or omitted during cDNA synthesis. (A) left gel: ORF9var_Fw – ORF9var_Rv: 1: unspliced, 2: splice variant 1, 3: splice variant 2, right gel: ORF9var_Fw – ORF9var1_spliceRv, ORF9var_Fw – ORF9var2_splice_Rv., (B) left gel: ORF48_Fw – ORF48_Rv: 1: unspliced, 2: spliced, right gel: ORF48_Fw – ORF48_spliceRv. (C) ORF12-13_Fw – ORF12-13_Rv: ORF12-13 fusion RNA. (D). ORF31-32_Fw – ORF31-32_Rv_#1 or ORF31-32_Fw – ORF31-32_Rv_#2. RNA31-32 fusion RNA (E) ORF13-5_Fw – ORF13-5_Rv: RNA13.5-1. (F) VZV_ORF43A_Fw – ORF43.5_Rv: RNA 43-2; 1: unspliced, 2: spliced.

**Figure S2. Related to Figure 2.**
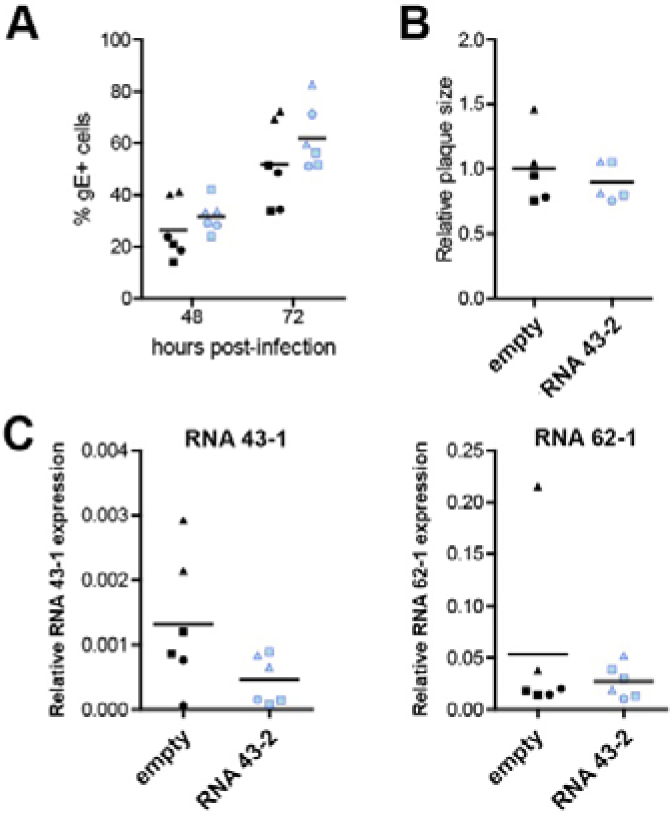
Functional analysis of VZV RNA 43-2. (A-C) ARPE-19 cell lines stably expressing empty pcDNA3.1 (black) or RNA 43-2 (light blue) were generated. Squares, circles and triangles represent the 3 independently generated stable ARPE-19 cell lines. (A-C) Cells were infected with cell-free VZV EMC-1. (A) Percentage of VZV-infected cells (glycoprotein E [gE]-positive) at 48 hpi or 72 hpi determined by flow cytometry. (B) Relative plaque size (normalized to empty pcDNA3.1 transfected cell lines) at 72 hpi. (C) Relative expression of RNA 43-1 and RNA 62-1 (normalized to β-actin transcripts) at 24 hpi.

**Figure S3.**
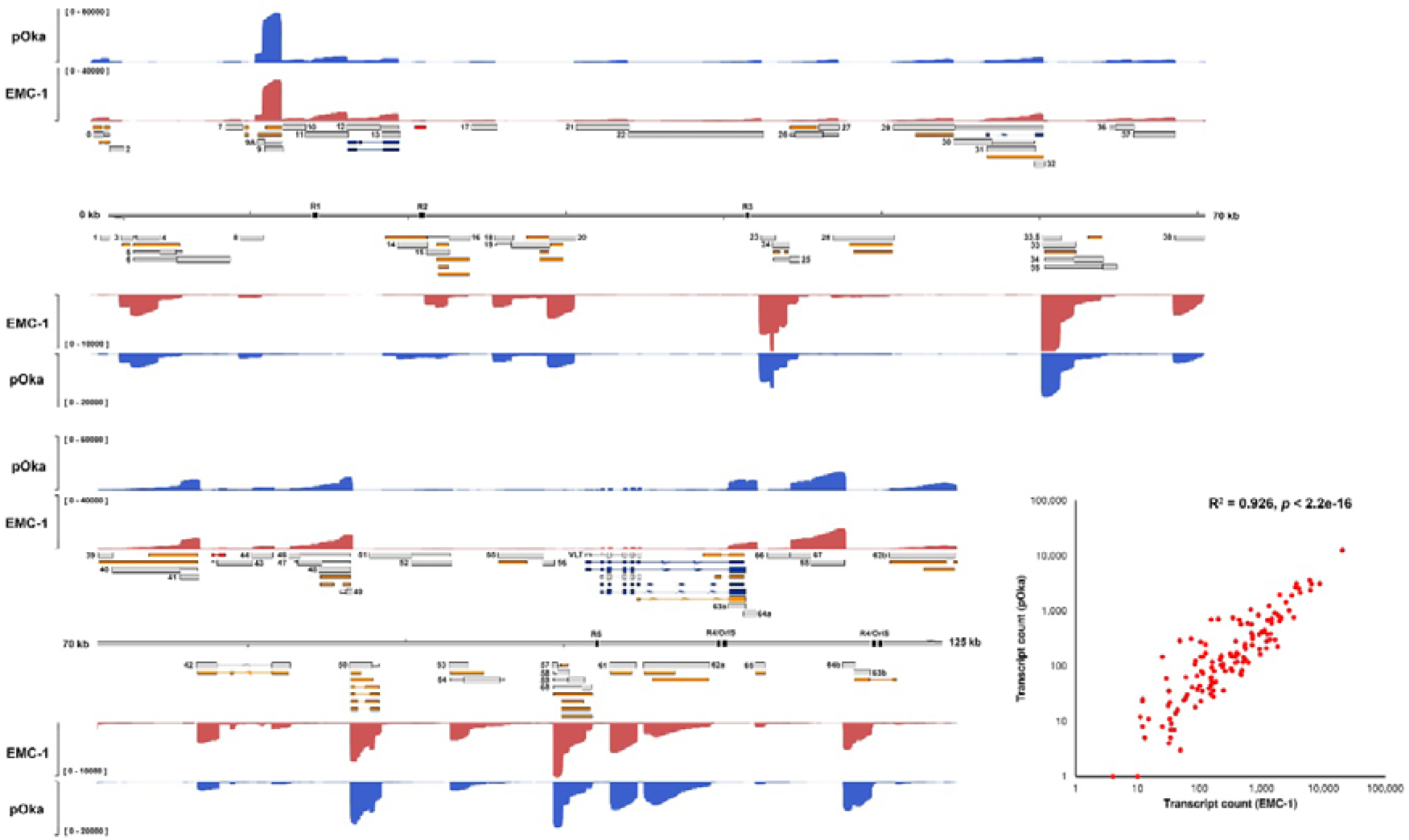
VZV transcriptome diversity is consistent between viral strains. Coverage plots derived from Nanopore dRNA-Seq of VZV strain pOka (blue) and VZV strain EMC-1 (red)-infected ARPE-19 cells at 96 hpi. Note that while relative and absolute abundances of distinct VZV RNAs differ between strains, all RNAs included in our annotation are represented. Y-axis denote coverage range. Canonical (grey), alternative (orange), fusion (dark blue) and putative ncRNAs (red) are shown with canonical CDS regions indicated by wide boxes and UTRs shown as thin boxes. The absence of a CDS region indicates that an RNA has uncertain coding potential. The reiterative repeat regions R1-R5 and both copies of the OriS are shown as black boxes embedded in the genome track. The scatter plot shows the correlation between VZV transcript abundance in VZV EMC-1-infected and VZV pOka-infected ARPE-19 cells. Pearson R^2^ and *p*-value are indicated.

**Figure S4. Related to Figure 3.**
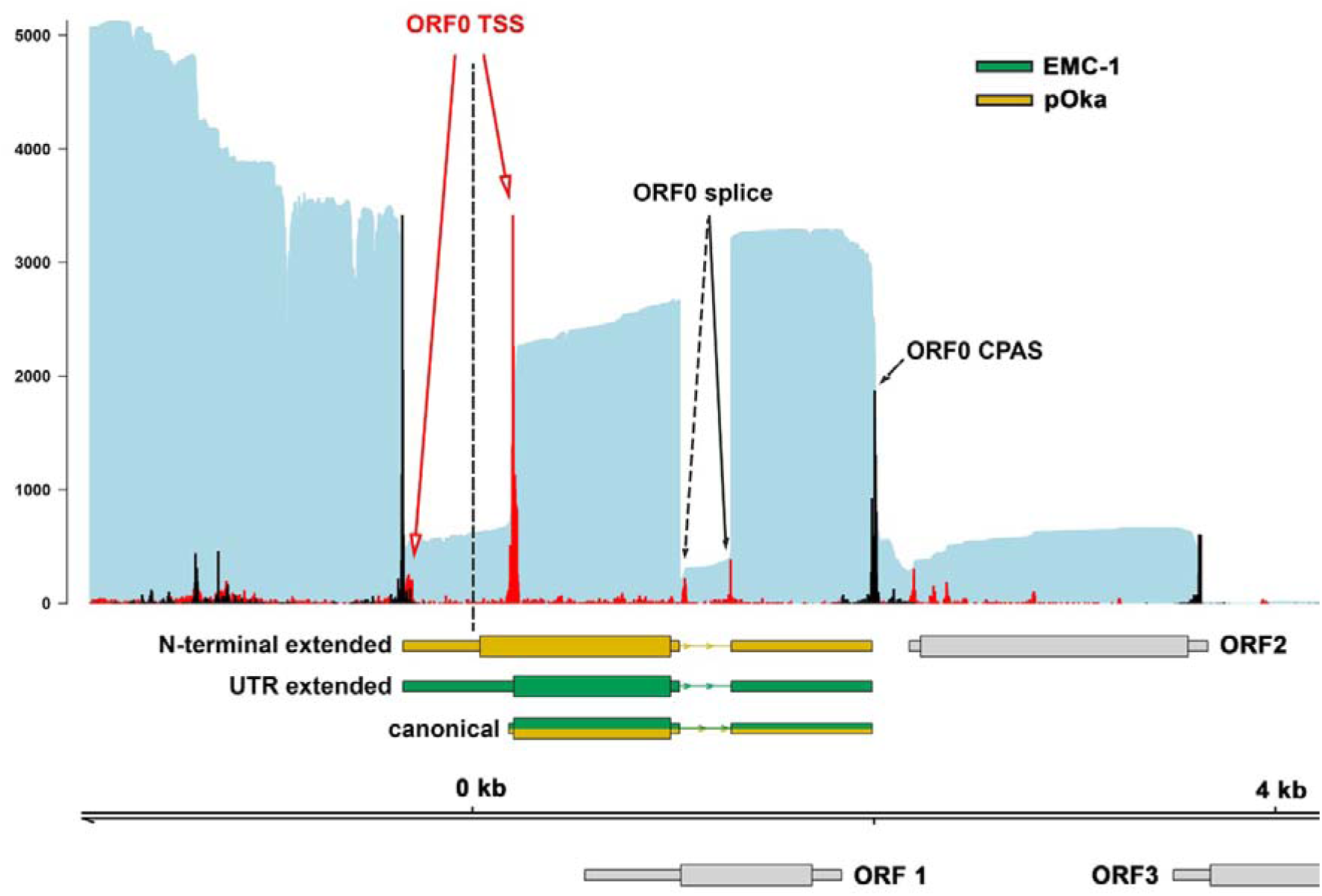
Expression of ORF0 RNA isoforms during lytic infection of ARPE-19 cells with VZV pOka and EMC-1. Identical TSS and CPAS sites were detected in ARPE-19 cells infected with cell-free VZV pOka (VZV clade 2; yellow) or -EMC-1 (VZV clade 1; green) at 96 hpi, yielding two ORF0 RNA isoforms. However, due to nucleotide variability between both strains, VZV pOka RNA 0-1 includes a larger CDS and can encode an alternative ORF0 protein variant, compared to VZV EMC-1. Nanopore dRNA-Seq (teal) coverage plots are shown. Y-axis values indicate the maximum read depth of that specific track. CDS regions and UTRs are shown as wide and thin boxes, respectively.

**Figure S5.**
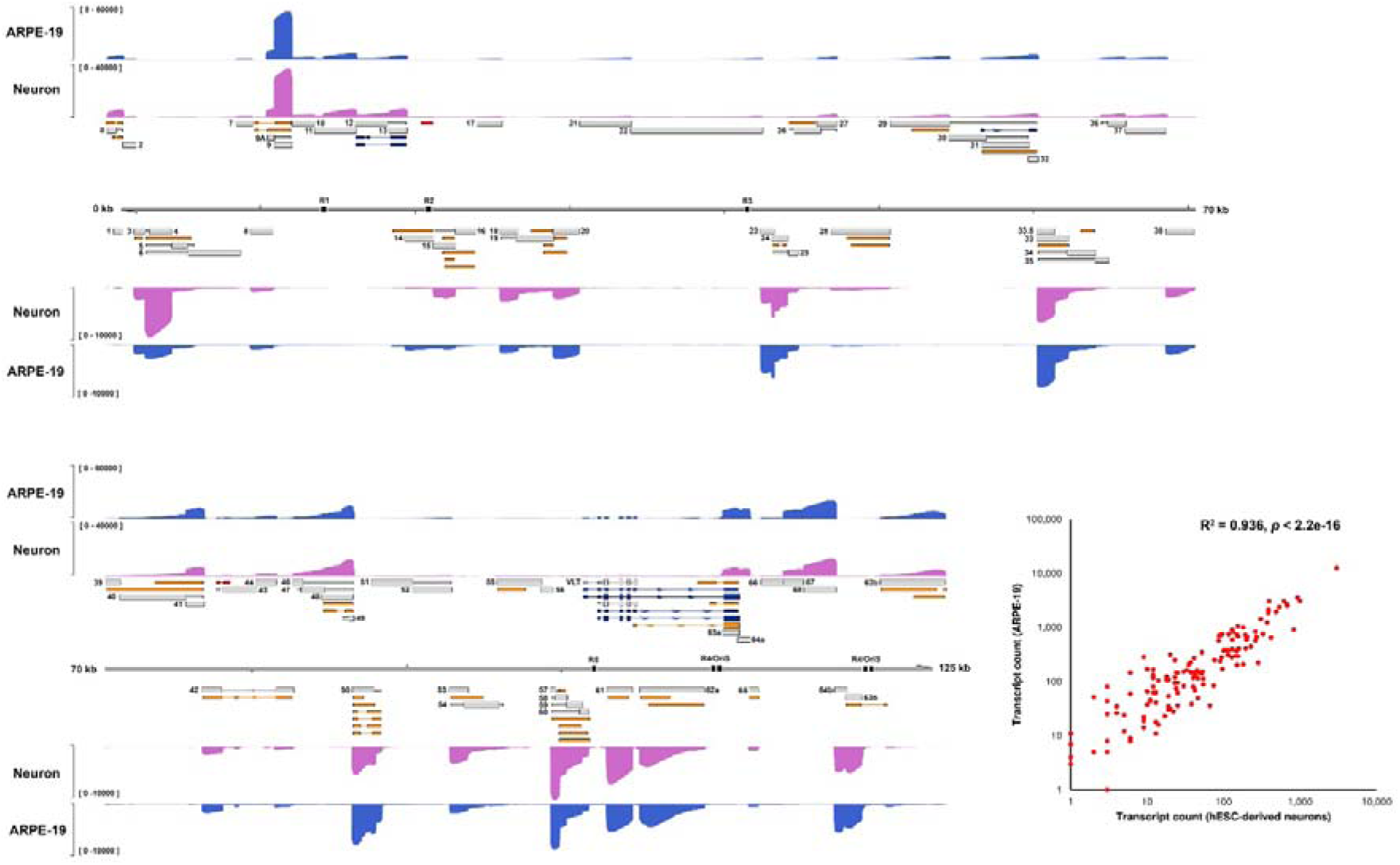
VZV transcriptome diversity is consistent between infected cell types. Coverage plots derived from Nanopore dRNA-Seq of VZV pOka infected ARPE-19 cells (blue) and hESC-derived neurons (purple). RNA was collected at 96 hpi (ARPE-19 cells) or 144 hpi (neurons). Note that while relative and absolute abundances of distinct VZV RNAs differ between cell types, all RNAs included in our annotation are represented. Y-axis denote coverage range. Canonical (grey), alternative (orange), fusion (dark blue) and putative ncRNAs (red) are shown with canonical CDS regions indicated by wide boxes and UTRs shown as thin boxes. The absence of CDS region indicates that the respective VZV RNA has an uncertain coding potential. Reiterative repeat regions R1-R5 and both copies of the OriS are shown as black boxes embedded in the genome track. The scatter plot shows the correlation between VZV transcript abundance in hESC-derived neurons and ARPE-19 cells. Pearson R2 and p-value are indicated.

**Figure S6. Related to Figure 4.**
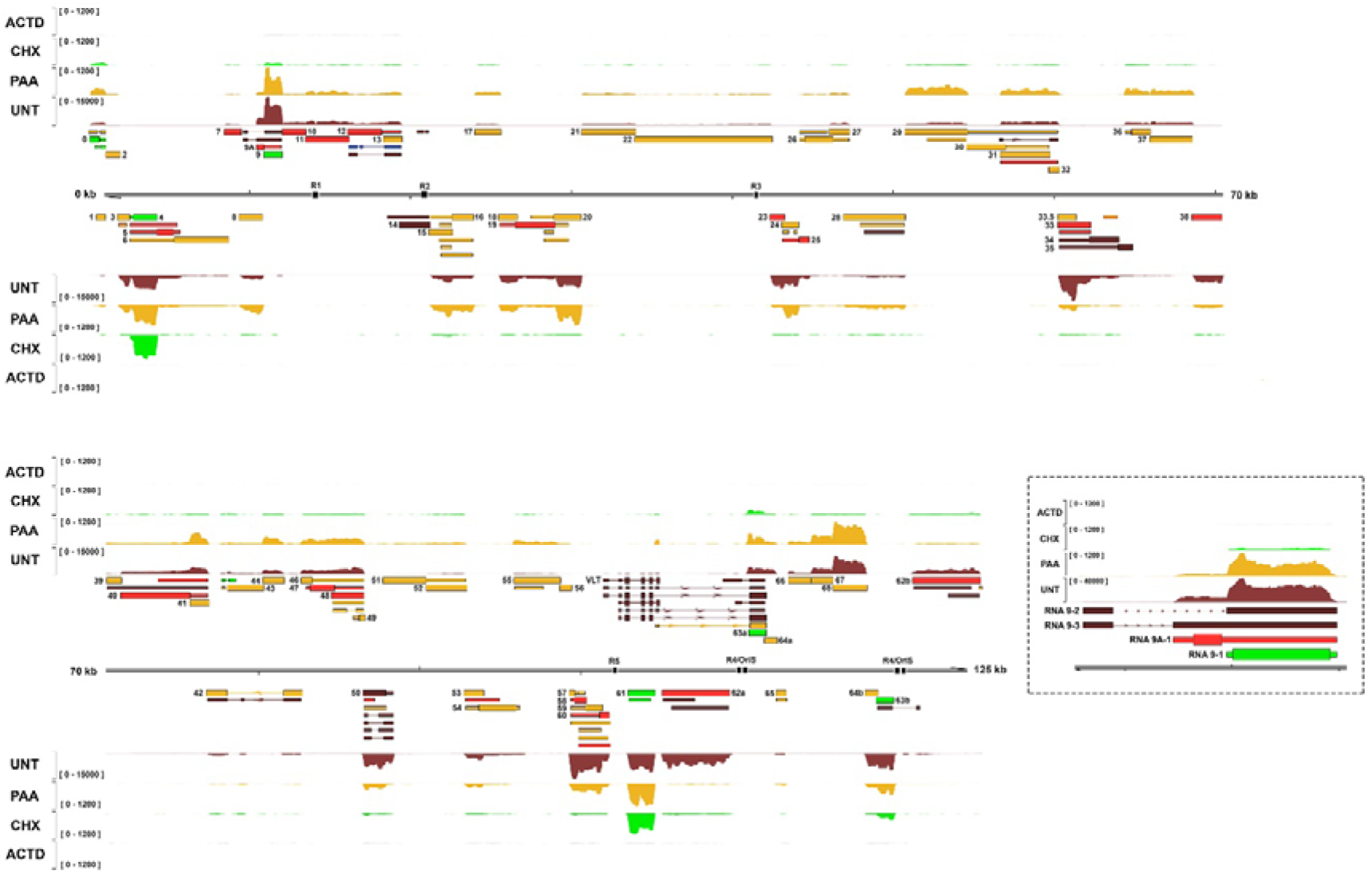
Kinetic class of viral RNAs expressed during lytic VZV infection of ARPE-19 cells. **(A)** Coverage plots derived from Illumina RNA-Seq of ARPE-19 cells infected with cell-free VZV EMC-1 and treated with actinomycin D (ActD; grey), cycloheximide (CHX; green), phosphonoacetic acid (PAA; gold), or untreated (UNT; red). Y-axis denotes coverage range. All transcripts are colour-coded according to their assigned kinetic class: *IE* – green, *E* – gold, *LL* – red, *TL* – dark red. Canonical CDS regions are indicated by wide boxes with UTRs shown as thin boxes. Absence of CDS region indicates that an RNA has an uncertain coding potential. Reiterative repeat regions R1-R5 and both copies of the OriS are shown as black boxes embedded in the genome track. (**B**) The inset black hatched box presents a close-up view of the RNA 9 locus.

**Figure S7. Related to Figure 6.**
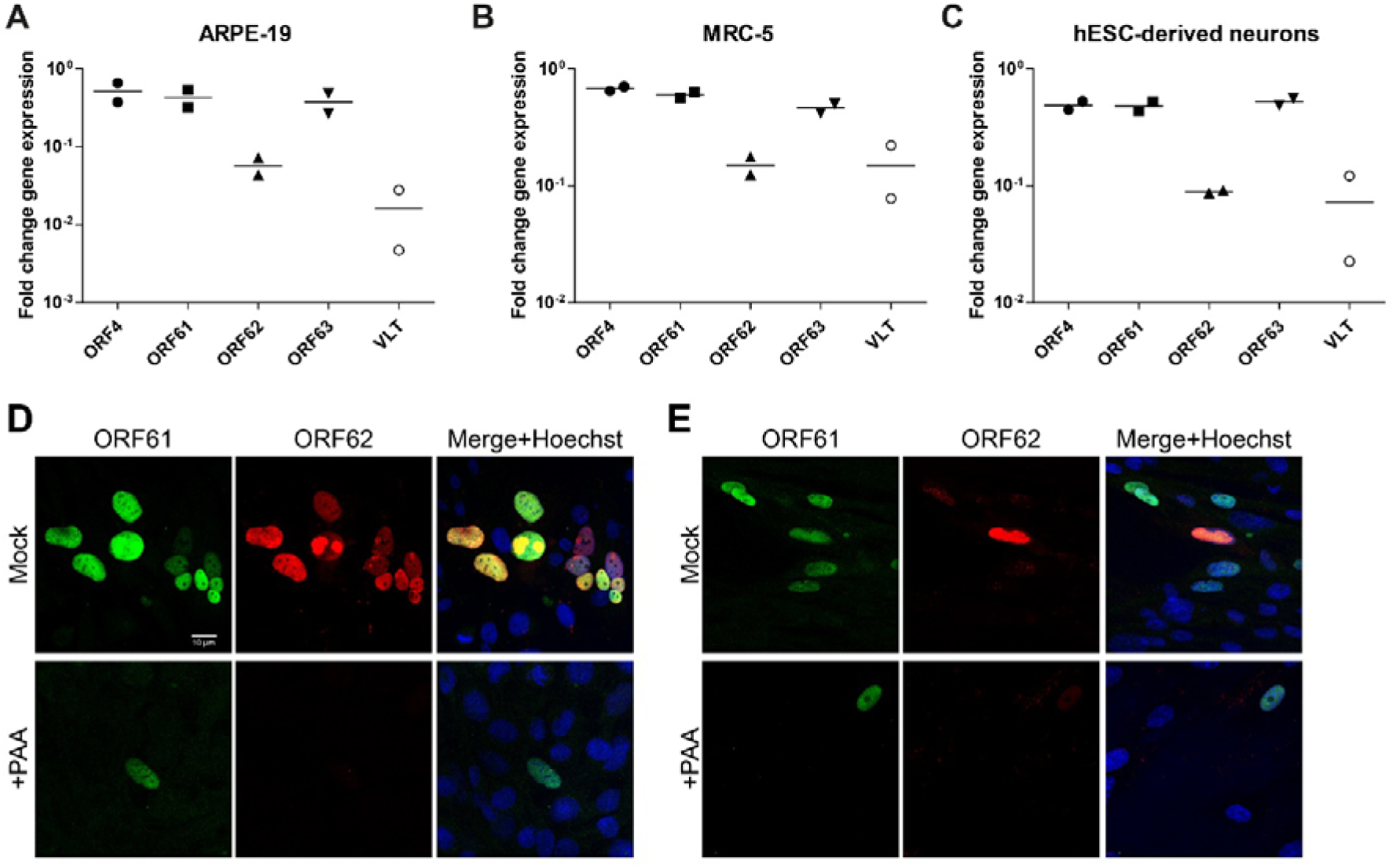
ORF62 transcripts are expressed with *Late* kinetics during VZV infection. (**A-C**) ARPE-19 cells, hESC-derived neurons and MRC-5 cells were infected with cell-free VZV strain pOka for 20-24 hours in the presence or absence of phosphonoformic acid. Fold change in gene expression in phosphonoformic acid-treated samples relative to untreated samples (calibrator) and normalized to β-actin using the 2^-ΔΔCt^ method. (**D-E**) Representative images of VZV strain EMC-1-infected MRC-5 (**D**) and MeWo (**E**) cells immunofluorescently stained for pORF61 (green) and pORF62 (red). Nuclei were counterstained with Hoechst (blue). Top row: 400x magnification, bottom rows: 1000x magnification.

**Figure S8.**
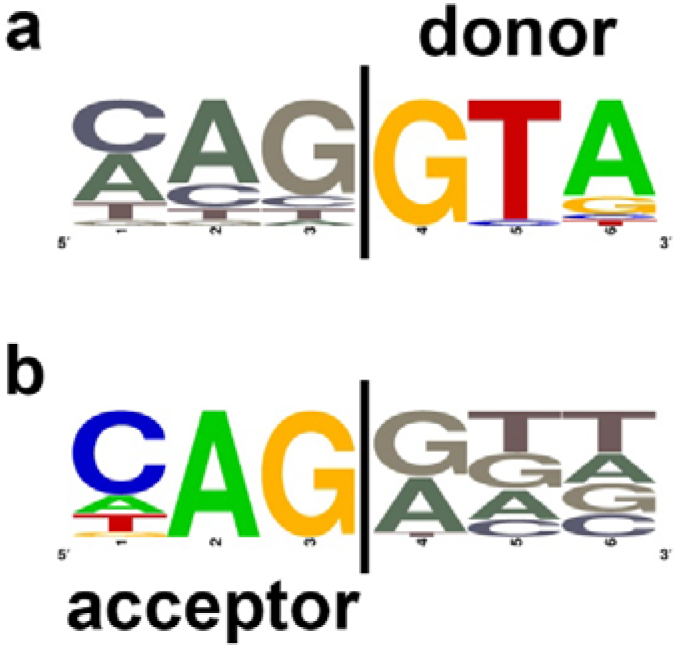
Splicing patterns in the VZV transcriptome. Twenty-eight VZV transcripts comprise two or more exons. The consensus **(A)** splice donor and **(B)** splice acceptor motifs were derived using WebLogo (Crooks et al., 2004).

## References

Arias, C., Weisburd, B., Stern-Ginossar, N., Mercier, A., Madrid, A.S., Bellare, P., Holdorf, M., Weissman, J.S., and Ganem, D. (2014). KSHV 2.0: A Comprehensive Annotation of the Kaposi’s Sarcoma-Associated Herpesvirus Genome Using Next-Generation Sequencing Reveals Novel Genomic and Functional Features. PLoS Pathog. 10.

Baird, N.L., Bowlin, J.L., Cohrs, R.J., Gilden, D., and Jones, K.L. (2014). Comparison of Varicella-Zoster Virus RNA Sequences in Human Neurons and Fibroblasts. J. Virol. 88, 5877–5880.

Bencun, M., Klinke, O., Hotz-Wagenblatt, A., Klaus, S., Tsai, M.H., Poirey, R., and Delecluse, H.J. (2018). Translational profiling of B cells infected with the Epstein-Barr virus reveals 5 leader ribosome recruitment through upstream open reading frames. Nucleic Acids Res. 46, 2802–2819.

Bonfert, T., Kirner, E., Csaba, G., Zimmer, R., and Friedel, C.C. (2015). ContextMap 2: Fast and accurate context-based RNA-seq mapping. BMC Bioinformatics 16.

Cohen, J.I. (2010). The Varicella-Zoster Virus Genome. A. Abendroth, A.M. Arvin, and J.F. Moffat, eds. (Berlin, Heidelberg: Springer Berlin Heidelberg), pp. 1–14.

Cox, E., Reddy, S., Iofin, I., and Cohen, J.I. (1998). Varicella-zoster virus ORF57, unlike its pseudorabies virus UL3.5 Homolog, is dispensable for viral replication in cell culture. Virology 250, 205–209.

Crooks, G.E., Hon, G., Chandonia, J.M., and Brenner, S.E. (2004). WebLogo: A sequence logo generator. Genome Res. 14, 1188–1190.

Davison, A.J., and Scott, J.E. (1986). The complete DNA sequence of varicella-zoster virus. J. Gen. Virol. 67, 1759–1816.

Defechereux, P., Melen, L., Baudoux, L., Merville-Louis, M.P., Rentier, B., and Piette, J. (1993). Characterization of the regulatory functions of varicella-zoster virus open reading frame 4 gene product. J. Virol. 67, 4379–4385.

Depledge, D.P., Sadaoka, T., and Ouwendijk, W.J.D. (2018a). Molecular aspects of varicella-zoster virus latency. Viruses 10.

Depledge, D.P., Ouwendijk, W.J.D., Sadaoka, T., Braspenning, S.E., Mori, Y., Cohrs, R.J., Verjans, G.M.G.M., and Breuer, J. (2018b). A spliced latency-associated VZV transcript maps antisense to the viral transactivator gene. Nat. Commun. 9.

Depledge, D.P., Mohr, I., and Wilson, A.C. (2018c). Going the Distance: Optimizing RNA-Seq Strategies for Transcriptomic Analysis of Complex Viral Genomes. J. Virol. 93.

Depledge, D.P., Srinivas, K.P., Sadaoka, T., Bready, D., Mori, Y., Placantonakis, D.G., Mohr, I., and Wilson, A.C. (2019). Direct RNA sequencing on nanopore arrays redefines the transcriptional complexity of a viral pathogen. Nat. Commun. 10.

Felser, J.M., Kinchington, P.R., Inchauspe, G., Straus, S.E., and Ostrove, J.M. (1988). Cell lines containing varicella-zoster virus open reading frame 62 and expressing the “IE” 175 protein complement ICP4 mutants of herpes simplex virus type 1. J. Virol. 62, 2076–2082.

Finkel, Y., Schmiedel, D., Tai-Schmiedel, J., Nachshon, A., Winkler, R., Dobesova, M., Schwartz, M., Mandelboim, O., and Stern-Ginossar, N. (2020). Comprehensive annotations of human herpesvirus 6A and 6B genomes reveal novel and conserved genomic features. Elife 9.

Garalde, D.R., Snell, E.A., Jachimowicz, D., Sipos, B., Lloyd, J.H., Bruce, M., Pantic, N., Admassu, T., James, P., Warland, A., et al. (2018). Highly parallel direct RNA sequencing on an array of nanopores. Nat. Methods 15, 201–206.

Gater, A., Uhart, M., McCool, R., and Préaud, E. (2015). The humanistic, economic and societal burden of Herpes Zoster in Europe: A critical review. BMC Public Health 15.

Gershon, A.A., Breuer, J., Cohen, J.I., Cohrs, R.J., Gershon, M.D., Gilden, D., Grose, C., Hambleton, S., Kennedy, P.G.E., Oxman, M.N., et al. (2015). Varicella zoster virus infection. Nat. Rev. Dis. Prim. 1, 15016.

Gilden, D.H., Vafai, A., Shtram, Y., Becker, Y., Devlin, M., and Wellish, M. (1983). Varicella-zoster virus DNA in human sensory ganglia. Nature 306, 478–480.

Gilden, D.H., Kleinschmidt-DeMasters, B.K., LaGuardia, J.J., Mahalingam, R., and Cohrs, R.J. (2000). Neurologic complications of the reactivation of varicella-zoster virus. N. Engl. J. Med. 342, 635–645.

Hahne, F., and Ivanek, R. (2016). Visualizing genomic data using Gviz and bioconductor. In Methods in Molecular Biology, pp. 335–351.

Hancock, M.H., and Skalsky, R.L. (2018). Roles of Non-coding RNAs During Herpesvirus Infection BT - Roles of Host Gene and Non-coding RNA Expression in Virus Infection. R.A. Tripp, and S.M. Tompkins, eds. (Cham: Springer International Publishing), pp. 243–280.

Heinz, S., Benner, C., Spann, N., Bertolino, E., Lin, Y.C., Laslo, P., Cheng, J.X., Murre, C., Singh, H., and Glass, C.K. (2010). Simple Combinations of Lineage-Determining Transcription Factors Prime cis-Regulatory Elements Required for Macrophage and B Cell Identities. Mol. Cell 38, 576–589.

Honess, R.W., and Roizman, B. (1974). Regulation of Herpesvirus Macromolecular Synthesis I. Cascade Regulation of the Synthesis of Three Groups of Viral Proteins 1. J. Virol. 14, 8–19.

Jensen, N.J., Depledge, D.P., Ng, T.F.F., Leung, J., Quinlivan, M., Radford, K.W., Folster, J., Tseng, H.-F., LaRussa, P., Jacobsen, S.J., et al. (2020). Analysis of the reiteration regions (R1 to R5) of varicella-zoster virus. Virology 546, 38–50.

Johnson, R.W., and Rice, A.S.C. (2014). Postherpetic neuralgia. N. Engl. J. Med. 371, 1526–1533.

Jones, M., Dry, I.R., Frampton, D., Singh, M., Kanda, R.K., Yee, M.B., Kellam, P., Hollinshead, M., Kinchington, P.R., O’Toole, E.A., et al. (2014). RNA-seq Analysis of Host and Viral Gene Expression Highlights Interaction between Varicella Zoster Virus and Keratinocyte Differentiation. PLoS Pathog. 10.

Kang, Y.J., Yang, D.C., Kong, L., Hou, M., Meng, Y.Q., Wei, L., and Gao, G. (2017). CPC2: A fast and accurate coding potential calculator based on sequence intrinsic features. Nucleic Acids Res. 45, W12–W16.

Kemble, G.W., Annunziato, P., Lungu, O., Winter, R.E., Cha, T.-A., Silverstein, S.J., and Spaete, R.R. (2000). Open Reading Frame S/L of Varicella-Zoster Virus Encodes a Cytoplasmic Protein Expressed in Infected Cells. J. Virol. 74, 11311 LP–11321.

Kinchington, P.R., Hougland, J.K., Arvin, A.M., Ruyechan, W.T., and Hay, J. (1992). The varicella-zoster virus immediate-early protein IE62 is a major component of virus particles. J. Virol. 66, 359–366.

Koshizuka, T., Ota, M., Yamanishi, K., and Mori, Y. (2010). Characterization of varicella-zoster virus-encoded ORF0 gene-Comparison of parental and vaccine strains. Virology 405, 280–288.

Kost, R.G., Kupinsky, H., and Straus, S.E. (1995). Varicella-Zoster Virus Gene 63: Transcript Mapping and Regulatory Activity. Virology 209, 218–224.

Lawrence, M., Huber, W., Pagès, H., Aboyoun, P., Carlson, M., Gentleman, R., Morgan, M.T., and Carey, V.J. (2013). Software for Computing and Annotating Genomic Ranges. PLoS Comput. Biol. 9.

Lenac Rovis, T., Bailer, S.M., Pothineni, V.R., Ouwendijk, W.J.D., Simic, H., Babic, M., Miklic, K., Malic, S., Verweij, M.C., Baiker, A., et al. (2013). Comprehensive Analysis of Varicella-Zoster Virus Proteins Using a New Monoclonal Antibody Collection. J. Virol. 87, 6943–6954.

Leppek, K., Das, R., and Barna, M. (2018). Functional 5’ UTR mRNA structures in eukaryotic translation regulation and how to find them. Nat. Rev. Mol. Cell Biol. 19, 158–174.

Li, H. (2018). Minimap2: Pairwise alignment for nucleotide sequences. Bioinformatics 34, 3094–3100.

Li, H., and Durbin, R. (2009). Fast and accurate short read alignment with Burrows-Wheeler transform. Bioinformatics 25, 1754–1760.

Li, H., Handsaker, B., Wysoker, A., Fennell, T., Ruan, J., Homer, N., Marth, G., Abecasis, G., and Durbin, R. (2009). The Sequence Alignment/Map format and SAMtools. Bioinformatics 25, 2078–2079.

Ma, H.T., and Poon, R.Y.C. (2016). Synchronization of HeLa cells. In Methods in Molecular Biology, pp. 189–201.

Maertzdorf, J., Remeijer, L., Van Der Lelij, A., Buitenwerf, J., Niesters, H.G.M., Osterhaus, A.D.M.E., and Verjans, G.M.G.M. (1999). Amplification of reiterated sequences of herpes simplex virus type 1 (HSV-1) genome to discriminate between clinical HSV-1 isolates. J. Clin. Microbiol. 37, 3518–3523.

Michael, E.J., Kuck, K.M., and Kinchington, P.R. (1998). Anatomy of the Varicella-Zoster Virus Open-Reading Frame 4 Promoter. J. Infect. Dis. 178, S27–S33.

Moriuchi, H., Moriuchi, M., Straus, S.E., and Cohen, J.I. (1993). Varicella-zoster virus (VZV) open reading frame 61 protein transactivates VZV gene promoters and enhances the infectivity of VZV DNA. J. Virol. 67, 4290–4295.

Moriuchi, M., Moriuchi, H., Straus, S.E., and Cohen, J.I. (1994). Varicella-Zoster Virus (VZV) Virion-Associated Transactivator Open Reading Frame 62 Protein Enhances the Infectivity of VZV DNA. Virology 200, 297–300.

Murata, M., Nishiyori-Sueki, H., Kojima-Ishiyama, M., Carninci, P., Hayashizaki, Y., and Itoh, M. (2014). Detecting expressed genes using CAGE. Methods Mol. Biol. 1164, 67–85.

Norberg, P., Depledge, D.P., Kundu, S., Atkinson, C., Brown, J., Haque, T., Hussaini, Y., MacMahon, E., Molyneaux, P., Papaevangelou, V., et al. (2015). Recombination of Globally Circulating Varicella-Zoster Virus. J. Virol. 89, 7133–7146.

O’Grady, T., Wang, X., Höner Zu Bentrup, K., Baddoo, M., Concha, M., and Flemington, E.K. (2016). Global transcript structure resolution of high gene density genomes through multi-platform data integration. Nucleic Acids Res. 44.

O’Grady, T., Feswick, A., Hoffman, B.A., Wang, Y., Medina, E.M., Kara, M., van Dyk, L.F., Flemington, E.K., and Tibbetts, S.A. (2019). Genome-wide Transcript Structure Resolution Reveals Abundant Alternate Isoform Usage from Murine Gammaherpesvirus 68. Cell Rep. 27, 3988–4002.e5.

Ouwendijk, W.J.D., Choe, A., Nagel, M.A., Gilden, D., Osterhaus, A.D.M.E., Cohrs, R.J., and Verjans, G.M.G.M. (2012). Restricted Varicella-Zoster Virus Transcription in Human Trigeminal Ganglia Obtained Soon after Death. J. Virol. 86, 10203–10206.

Ouwendijk, W.J.D., Mahalingam, R., de Swart, R.L., Haagmans, B.L., van Amerongen, G., Getu, S., Gilden, D., Osterhaus, A.D.M.E., and Verjans, G.M.G.M. (2013). T-Cell Tropism of Simian Varicella Virus during Primary Infection. PLoS Pathog. 9.

Ouwendijk, W.J.D., Geluk, A., Smits, S.L., Getu, S., Osterhaus, A.D.M.E., and Verjans, G.M.G.M. (2014). Functional Characterization of Ocular-Derived Human Alphaherpesvirus Cross-Reactive CD4 T Cells. J. Immunol. 192, 3730 LP–3739.

Ouwendijk, W.J.D., Dekker, L., Van den Ham, H.-J., Lenac Roviš, T., Haefner, E.S., Jonjic, S., Haas, J., Luider, T.M., and Verjans, G.M.G.M. (2020). Analysis of virus and host proteomes during productive HSV-1 and VZV infection in human epithelial cells. Front. Microbiol.

Pelechano, V., and Steinmetz, L.M. (2013). Gene regulation by antisense transcription. Nat. Rev. Genet. 14, 880–893.

Perera, L.P., Mosca, J.D., Sadeghi-Zadeh, M., Ruyechan, W.T., and Hay, J. (1992). The Varicella-Zoster virus immediate early protein, IE62, can positively regulate its cognate promoter. Virology 191, 346–354.

Perera, L.P., Mosca, J.D., Ruyechan, W.T., Hayward, G.S., Straus, S.E., and Hay, J. (1993). A major transactivator of varicella-zoster virus, the immediate-early protein IE62, contains a potent N-terminal activation domain. J. Virol. 67, 4474–4483.

Prazsák, I., Moldován, N., Balázs, Z., Tombácz, D., Megyeri, K., Szucs, A., Csabai, Z., and Boldogkoi, Z. (2018). Long-read sequencing uncovers a complex transcriptome topology in varicella zoster virus. BMC Genomics 19.

Quinlan, A.R., and Hall, I.M. (2010). BEDTools: A flexible suite of utilities for comparing genomic features. Bioinformatics 26, 841–842.

Reichelt, M., Brady, J., and Arvin, A.M. (2009). The Replication Cycle of Varicella-Zoster Virus: Analysis of the Kinetics of Viral Protein Expression, Genome Synthesis, and Virion Assembly at the Single-Cell Level. J. Virol. 83, 3904–3918.

Ruyechan, W.T., Peng, H., Yang, M., and Hay, J. (2003). Cellular factors and IE62 activation of VZV promoters. J. Med. Virol. 70.

Sadaoka, T., Yoshii, H., Imazawa, T., Yamanishi, K., and Mori, Y. (2007). Deletion in Open Reading Frame 49 of Varicella-Zoster Virus Reduces Virus Growth in Human Malignant Melanoma Cells but Not in Human Embryonic Fibroblasts. J. Virol. 81, 12654–12665.

Sadaoka, T., Yanagi, T., Yamanishi, K., and Mori, Y. (2010). Characterization of the Varicella-Zoster Virus ORF50 Gene, Which Encodes Glycoprotein M. J. Virol. 84, 3488–3502.

Sadaoka, T., Depledge, D.P., Rajbhandari, L., Venkatesan, A., Breuer, J., and Cohen, J.I. (2016). In vitro system using human neurons demonstrates that varicella-zoster vaccine virus is impaired for reactivation, but not latency. Proc. Natl. Acad. Sci. 113, E2403–E2412.

Sadaoka, T., Schwartz, C.L., Rajbhandari, L., Venkatesan, A., and Cohen, J.I. (2017). Human Embryonic Stem Cell-Derived Neurons Are Highly Permissive for Varicella-Zoster Virus Lytic Infection. J. Virol. 92.

Sadaoka, T., Rajbhandari, L., Shukla, P., Jagdish, B., Lee, H., Lee, G., and Venkatesan, A. (2020). Human stem cell derived sensory neurons are positioned to support varicella zoster virus latency. BioRxiv 2020.01.24.919290.

Saxena, A., and Carninci, P. (2011). Long non-coding RNA modifies chromatin: Epigenetic silencing by long non-coding RNAs. BioEssays 33, 830–839.

Stern-Ginossar, N., Weisburd, B., Michalski, A., Le, V.T.K., Hein, M.Y., Huang, S.X., Ma, M., Shen, B., Qian, S.B., Hengel, H., et al. (2012). Decoding human cytomegalovirus. Science (80-.). 338, 1088–1093.

Storlie, J., Jackson, W., Hutchinson, J., and Grose, C. (2006). Delayed Biosynthesis of Varicella-Zoster Virus Glycoprotein C: Upregulation by Hexamethylene Bisacetamide and Retinoic Acid Treatment of Infected Cells. J. Virol. 80, 9544–9556.

Tang, A.D., Soulette, C.M., van Baren, M.J., Hart, K., Hrabeta-Robinson, E., Wu, C.J., and Brooks, A.N. (2020). Full-length transcript characterization of SF3B1 mutation in chronic lymphocytic leukemia reveals downregulation of retained introns. Nat. Commun. 11.

Thorvaldsdóttir, H., Robinson, J.T., and Mesirov, J.P. (2013). Integrative Genomics Viewer (IGV): High-performance genomics data visualization and exploration. Brief. Bioinform. 14, 178–192.

Toropova, K., Huffman, J.B., Homa, F.L., and Conway, J.F. (2011). The Herpes Simplex Virus 1 UL17 Protein Is the Second Constituent of the Capsid Vertex-Specific Component Required for DNA Packaging and Retention. J. Virol. 85, 7513–7522.

Tyler, S.D., Peters, G.A., Grose, C., Severini, A., Gray, M.J., Upton, C., and Tipples, G.A. (2007). Genomic cartography of varicella-zoster virus: A complete genome-based analysis of strain variability with implications for attenuation and phenotypic differences. Virology 359, 447–458.

Wang, L., Sommer, M., Rajamani, J., and Arvin, A.M. (2009). Regulation of the ORF61 Promoter and ORF61 Functions in Varicella-Zoster Virus Replication and Pathogenesis. J. Virol. 83, 7560–7572.

Whisnant, A.W., Jürges, C.S., Hennig, T., Wyler, E., Prusty, B., Rutkowski, A.J., L’hernault, A., Göbel, M., Döring, K., Menegatti, J., et al. (2019). Integrative functional genomics decodes herpes simplex virus 1. BioRxiv 603654.

Yang, M., Hay, J., and Ruyechan, W.T. (2004). The DNA Element Controlling Expression of the Varicella-Zoster Virus Open Reading Frame 28 and 29 Genes Consists of Two Divergent Unidirectional Promoters Which Have a Common USF Site. J. Virol. 78, 10939–10952.

Yang, M., Peng, H., Hay, J., and Ruyechan, W.T. (2006). Promoter Activation by the Varicella-Zoster Virus Major Transactivator IE62 and the Cellular Transcription Factor USF. J. Virol. 80, 7339–7353.

Zhang, C., Hastings, M.L., Krainer, A.R., and Zhang, M.Q. (2007). Dual-specificity splice sites function alternatively as 5’ and 3’ splice sites. Proc. Natl. Acad. Sci. U. S. A. 104, 15028–15033.

Zhang, Z., Selariu, A., Warden, C., Huang, G., Huang, Y., Zaccheus, O., Cheng, T., Xia, N., and Zhu, H. (2010). Genome-wide mutagenesis reveals that ORF7 is a novel VZV skin-tropic factor. PLoS Pathog. 6, 1–9.

